# The atypical Rho GTPase Rnd2 is critical for dentate granule neuron development and anxiety-like behavior during adult but not neonatal neurogenesis

**DOI:** 10.1101/2020.09.10.290866

**Authors:** Thomas Kerloch, Fanny Farrugia, Marlène Maître, Geoffrey Terral, Muriel Koehl, Julian Ik-Tsen Heng, Mylène Blanchard, Hélène Doat, Thierry Leste-Lasserre, Adeline Goron, Delphine Gonzales, François Guillemot, Djoher Nora Abrous, Emilie Pacary

## Abstract

Despite the central role of Rho GTPases in neuronal development, their functions in adult hippocampal neurogenesis remain poorly explored. Here, by using a retrovirus-based loss-of-function approach *in vivo*, we show that the atypical Rho GTPase Rnd2 is crucial for the survival, positioning, somatodendritic morphogenesis and functional maturation of adult-born dentate granule neurons. Interestingly, most of these functions are specific to granule neurons generated during adulthood since the deletion of *Rnd2* in neonatally-born granule neurons only affects dendritogenesis. In addition, suppression of *Rnd2* in adult-born dentate granule neurons increases anxiety-like behaviour whereas its deletion in pups has no such effect, a finding supporting the adult neurogenesis hypothesis of anxiety disorders. Thus, our results provide mechanistic insight into the differential regulation of hippocampal neurogenesis during development and adulthood, and establishes a causal relationship between Rnd2 expression and anxiety.

## INTRODUCTION

For decades, neurogenesis was believed to be restricted to embryonic and early postnatal periods in the mammalian brain. However, over the last 20 years, research has firmly established that new neurons are continuously born throughout the lifespan of mammals, especially in the hippocampal dentate gyrus (DG) (Goncalves et al., 2016). Indeed, although the majority of dentate granule neurons (DGNs) are generated in early postnatal life, new DGNs continue to be produced throughout adulthood in mammals albeit at lower rates (Altman and Bayer, 1990; Schlessinger et al., 1975), including in humans (Boldrini et al., 2018; Eriksson et al., 1998; Moreno-Jiménez et al., 2019; Spalding et al., 2013). Mammalian adult hippocampal neurogenesis (AHN) is a highly regulated, activity-dependent process that is reported to be critically involved in hippocampus-dependent functions such as memory encoding and mood regulation (Anacker and Hen, 2017).

Comparable to embryonic neurodevelopment, AHN is a multi-step process that begins with a phase of proliferation of neuronal progenitors (Abrous et al., 2005). Once generated, immature neurons migrate from the subgranular zone (SGZ) of the DG to the inner granule cell layer (GCL) and differentiate into glutamatergic DGNs. Newborn neurons then extend dendrites toward the molecular layer, project axons through the hilus toward the CA3 (Sun et al., 2013) and finally integrate into the preexisting neural circuitry. However only a few newborn cells are incorporated into the DG circuitry since the majority of these cells undergo apoptosis at the immature neuron stage (Kempermann et al., 2003; Tashiro et al., 2006a).

The finding that new neurons are generated throughout life has significant implications for brain repair. This has led to tremendous efforts to characterize how new neurons differentiate and integrate into adult neural circuitry. Yet, the cellular and molecular mechanisms that regulate the neurogenic process in the adult brain are not well characterized (Toda et al., 2018). Furthermore, the mechanisms that differently regulate adult versus developmental neurogenesis are poorly understood. Thus, further studies are needed to provide new insights into the mechanisms involved in the generation, survival and integration of individual newborn neurons in the adult versus developing brain. Such investigations might support the identification of new putative therapeutic targets for stimulating neurogenesis in health and disease.

Over the past several years, it has become clear that the Rho GTPases, the master regulators of the cytoskeleton, play a central role in various aspects of neuronal development, including migration, dendrite outgrowth, spine formation and maintenance (Govek et al., 2011; Govek et al., 2005). Yet, their functions during adult neurogenesis remain largely unexplored (Vadodaria and Jessberger, 2013). In particular, in the context of AHN, only few studies have addressed the roles played by the most well-characterized members of this family, Cdc42, Rac1 and RhoA. Vadodaria et al (2013) showed that Cdc42 is involved in mouse neural stem/progenitor cell proliferation, dendritic development and spine maturation, while Rac1 is important in the late steps of dendritic growth and spine maturation in adult hippocampal newborn neurons (Vadodaria et al., 2013). In the case of RhoA, pharmacological blockade of its signaling *in vivo* is reported to promote the survival of adult-born DGNs (Christie et al., 2013).

In this study, we have focused our attention on Rnd2 (also called Rho7/RhoN). Rnd2 belongs to the Rnd subfamily of atypical Rho proteins that lack intrinsic GTPase activity and are therefore constitutively bound to GTP (Chardin, 2006). Interestingly, among the 23 members of the Rho GTPase family (Grise et al., 2009), only *Rnd2* (and to a lesser extent *TC10*) is selectively enriched in the adult mouse SGZ and inner GCL, suggesting a potential prominent role in AHN (Miller et al., 2013). In addition, Rnd2 regulates embryonic cortical neurogenesis especially radial migration through RhoA inhibition (Heng et al., 2008; Pacary et al., 2011) and was shown to control multiple aspects of neuronal development *in vitro* like neurite outgrowth and branching (Azzarelli et al., 2015; Fujita et al., 2002; Tanaka et al., 2006). Thus, in view of these data, we postulated that Rnd2 might be a good candidate to play a key role in the regulation of AHN. Here, to elucidate the *in vivo* function of Rnd2 in this process, we used a retrovirus-based loss-of-function approach. With this strategy, we demonstrate that Rnd2 is crucial for the survival of adult-born DGNs but also for their correct positioning in the GCL, the morphogenesis of their somatodendritic compartment as well as their intrinsic excitability. In addition, we show that the expression of Rnd2 in adult-born DGNs mediates anxiety-like behaviour, indicating that Rnd2 cell autonomously influences the development of adult-born DGNs and controls anxiety behavior from these cells. Furthermore, we find that retrovirally-transduced Cre-mediated loss of *Rnd2* expression in neonatally-born (P0) DGNs only impacts dendrite formation, suggesting that Rnd2 plays distinct roles during developmental and adult neurogenesis in the DG.

## RESULTS

### Rnd2 is enriched in the temporal SGZ of the adult mouse DG

We began this study by examining the expression of Rnd2 in the mouse DG. We first observed by RNA *in situ* hybridization and real-time PCR that *Rnd2* expression decreases significantly in the DG between the postnatal and the adult period (Fig. 1A-D), similarly to related family members *Rnd1* and *Rnd3* (Fig. S1A, B). However in the adult brain, *Rnd2* shows a different pattern of expression compared to other members of the Rnd family, particularly interesting in the context of adult neurogenesis. Indeed, *Rnd2* is restricted to the SGZ and inner GCL of the DG (Fig. 1B, C), whereas *Rnd1* and *Rnd3* are more prominently localized to the CA1-CA3 hippocampal fields and in the DG (Fig. S1C). In addition, whereas *Rnd1* and *Rnd3* levels do not fluctuate dramatically along the septo-temporal axis (Fig. S1D), *Rnd2* mRNA expression increases in the temporal part of the adult DG (Fig. 1E; F_4,16_ = 7.16, p=0.002). This result is consistent with a RNA-seq study (http://hipposeq.janelia.org), which reported a significantly higher level of *Rnd2* mRNA expression in the temporal DGNs relative to septal DGNs (Cembrowski et al., 2016). By immunohistochemistry, we confirmed, at the protein level, the enrichment of Rnd2 in the SGZ and inner GCL of the adult DG (Fig. 1F) and we showed that Rnd2 is expressed in adult newborn neurons, by colabeling with doublecortin (DCX), a marker of immature neurons (Fig. 1G-I). The punctate immunostaining pattern for Rnd2 in these hippocampal cells is consistent with previous reports describing its localization to endosomes (Pacary et al., 2011; Tanaka et al., 2002) (Fig. 1F-I). Altogether these results indicate that Rnd2 is particularly enriched in the SGZ of the adult DG where newborn neurons are generated, suggesting that this atypical Rho GTPase might be involved in AHN.

**Figure 1:**
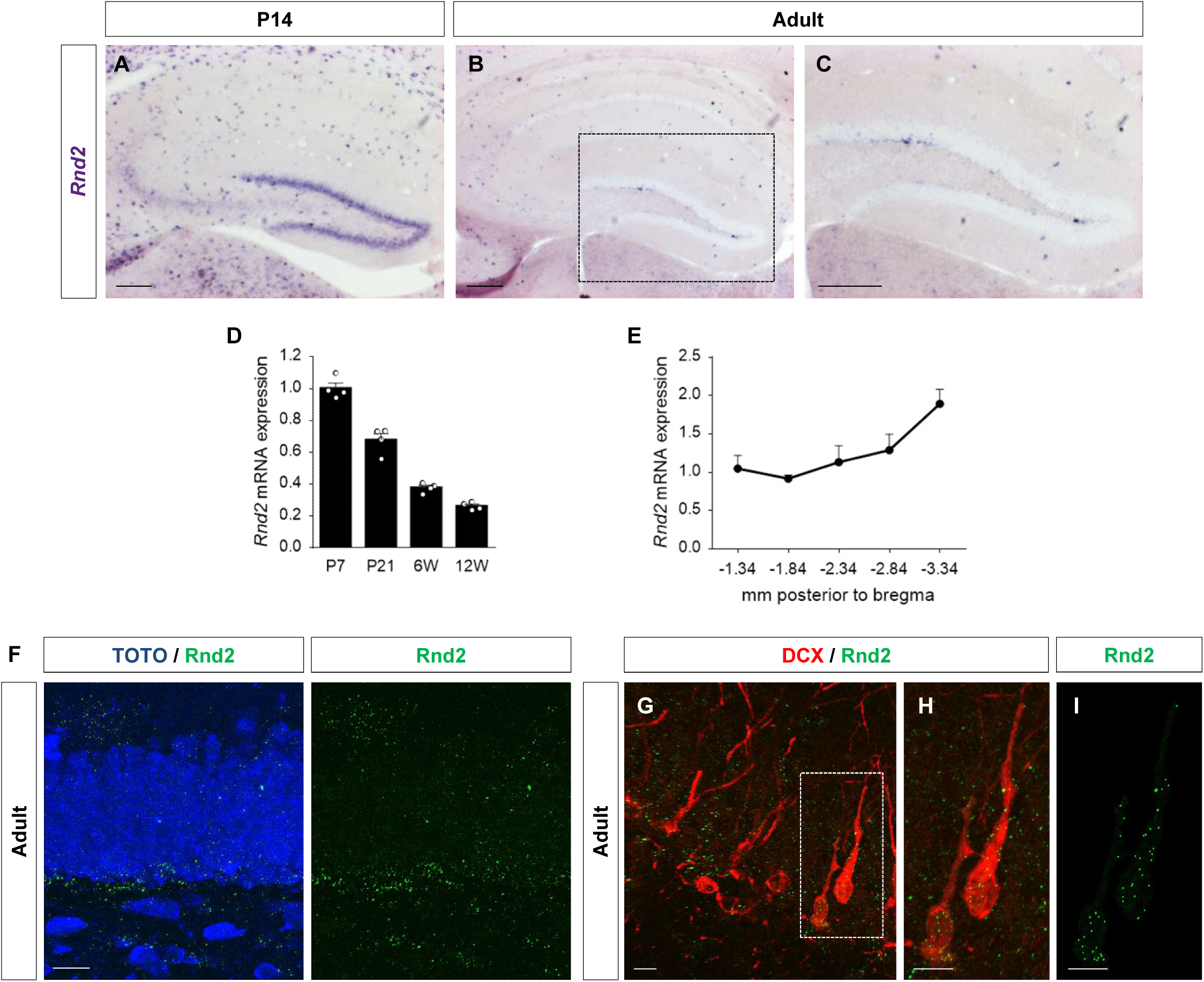
Rnd2 expression in the mouse DG. (**A, B, C**) Distribution of *Rnd2* transcripts in the hippocampus at postnatal day 14 (P14) (A) and at adult stage (12-week-old mouse) (B, C). The black rectangle in B shows the area enlarged in the inset C. (**D**) Analysis by real-time PCR of *Rnd2* mRNA expression in microdissected DG (SGZ and GCL) at different ages. Data are presented as fold change compared with the expression level at postnatal day 7 (P7) ± s.e.m (n = 4 mice per time point). P, postnatal day; W, weeks. (**E**) Analysis by real-time PCR of *Rnd2* mRNA expression along the septo-temporal axis in the adult (12-week-old) microdissected DG. Data are presented as fold change compared with the expression level at the anteroposterior coordinate −1.34 from the Bregma ± s.e.m (n = 5 mice per time point). (**F**) Adult (12-week-old) coronal sections immunostained for Rnd2. Nuclei were labeled with TOTO. (**G, H**) Immunostaining for Rnd2 and doublecortin (DCX) in the DG of a 12-week-old mouse. The white rectangle in G shows the area enlarged in the insets H and I. Imaris software was used to apply masks and selects the Rnd2 staining (in green) that was only inside DCX+ neurons. (**I**) Dots illustrating Rnd2 positive vesicles inside DCX+ neurons after Imaris analysis. Scale bars represent 200 µm (A, B, C), 20 µm (F) and 10 µm (G, H, I).

### Rnd2 is critical for the survival of adult-born DGNs

To address the role of Rnd2 in AHN, we deleted *Rnd2* specifically in adult-born DGNs using a loss of function approach based on retrovirus-mediated single-cell gene knockout in new neurons *in vivo* (Tashiro et al., 2006b). The principle of the technique is to deliver the Cre recombinase selectively to newborn neurons, using a retroviral vector, into the DG of *Rnd2*^*flox/flox*^ adult mice (Fig. S2A). By injecting retroviruses expressing Cre fused to green fluorescent protein (GFP/Cre, *Rnd2* deletion), or GFP only in control condition, into the DG of adult *Rnd2*^*flox/flox*^ mice (12-week-old), we first confirmed, after microdissection of GFP+ cells followed by RT-PCR, that *Rnd2* mRNA expression was strongly decreased in the GFP/Cre group compared to control whereas *Rnd1* and *Rnd3* mRNA expressions were not changed (Fig. S2B; t_15_=4.42, p<0.001 for Rnd2 analysis). Indeed, 7 days post-injection (dpi), the expression of *Rnd2* was already decreased by almost 80% (Fig. S2B).

Once validated, we used this viral strategy to study the role of Rnd2 in the development of adult-born DGNs. We first showed that *Rnd2* deletion in these cells does not affect neuronal progenitor proliferation and neuronal differentiation (Fig. S3). Intriguingly, we noticed that the number of GFP+ labeled cells in *Rnd2*-deleted animals was consistently lower compared to control animals 2-3 weeks after retroviral injection (Fig. 2A). In order to understand if Rnd2 expression is relevant to the survival of adult hippocampal newborn neurons, we quantified the number of GFP+ cells in the DG of *Rnd2*^*flox/flox*^ mice, injected with Cre/GFP or GFP expressing retrovirus, across multiple time points post-injection (Fig. 2B). Importantly, for each type of virus, the same preparation was used for all time point analyzed. In control condition, the number of retrovirus-labeled GFP cells at 21 dpi was reduced by 64% compared to 3 dpi (Fig. 2B), a finding which is consistent with previous studies (Kempermann et al., 2003; Tashiro et al., 2006a). Consistent with our first observations, the suppression of *Rnd2* substantially decreased the number of newborn DGNs from 14 dpi and, at 21 dpi, only 10.9% ± 1.9% of *Rnd2*-deleted cells survived compared to 35.7 ± 5.7% of control cells (Fig. 2A, B; t_9_=2.54, p=0.03 at 14 dpi; t_8_=4.12, p=0.003 at 21 dpi).

**Figure 2:**
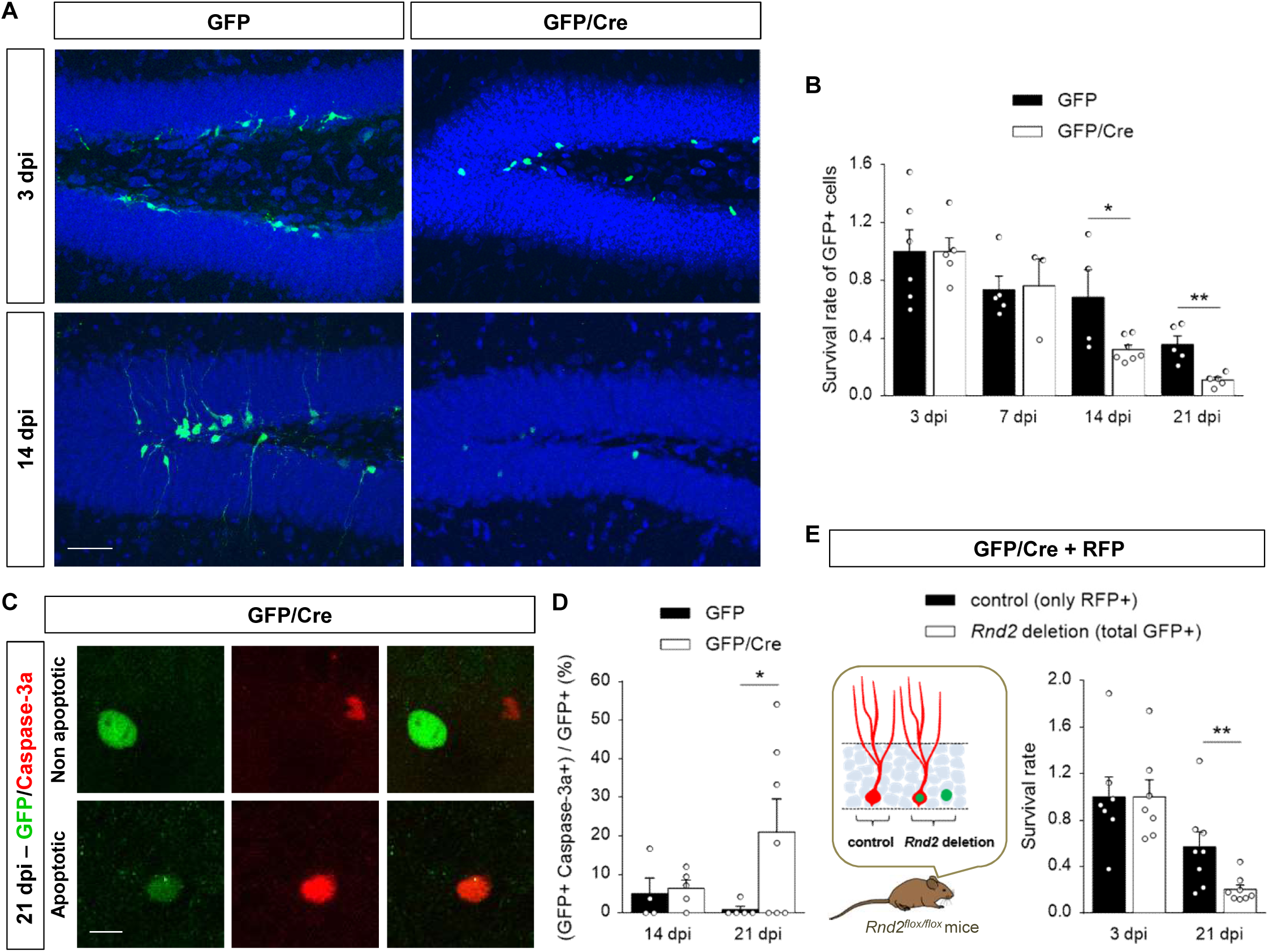
The absence of *Rnd2* reduces the survival of adult-born DGNs. (**A**)Images of the DG illustrating the reduction of GFP+ cell number after *Rnd2* deletion (GFP/Cre panel) compared to control (GFP panel) at 14 days post-injection (dpi). No difference was observed at 3 dpi. Nuclei were labeled with TOTO. (**B**) Survival rate of GFP+ cells at different time points. Mean ± s.e.m.; unpaired two-tailed Student’s t-test; *p < 0.05, **p < 0.01 (n = 3-7 mice). (**C**) Immunostaining for GFP and activated caspase 3 (caspase-3a) in the DG, 21 days after GFP/Cre virus injection. (**D**) Quantification of the percentage of transduced cells that are caspase-3a+ at 14 and 21 dpi. Mean ± s.e.m.; unpaired one-tailed Student’s t-test; *p < 0.05 (n = 4-7 mice). (**E**) The retroviral co-injection strategy allows to analyze in the same mice *Rnd2*-deleted and control new neurons. The graph shows the survival rate of control (RFP+ only) and *Rnd2*-knockout (GFP+ only and GFP+/RFP+) new neurons in *Rnd2*^*flox/flox*^ mice. Mean ± s.e.m.; paired two-tailed Student’s t-test; **p < 0.01 (n = 7-8 mice). Scale bars represent 50 µm (A) and 5 µm (C).

To determine whether the decrease in the number of new neurons after *Rnd2* suppression was due to their elimination through apoptosis, we performed immunostaining for activated caspase 3 (caspase 3a, Fig. 2C). Since apoptotic cells are rapidly cleared (Savill, 1997; Sierra et al., 2010), the probability of detecting apoptotic newborn cells is low. This is reflected in our observation that only a small fraction of GFP+ cells were immunopositive for caspase 3a in the control group at 14 and 21 dpi (Fig. 2D). Nonetheless, the proportion of transduced cells expressing caspase-3a was significantly increased at 21 dpi in the GFP/Cre condition compared to GFP (Fig. 2D; t_10_=1.98, p=0.04) indicating that *Rnd2* suppression in newborn neurons leads to enhanced programmed cell death.

To further confirm these results, we used a retroviral co-injection strategy (Tashiro et al., 2006a) in which we injected *Rnd2*^*flox/flox*^ mice with a retrovirus expressing GFP/Cre together with a red fluorescent protein (RFP) expressing control virus. This approach enables us to investigate viability of *Rnd2*-knockout (GFP+/RFP+ cells and GFP+ only) and control new neurons (RFP+ only) in the same mice (Fig. 2E). Consistent with our previous results, we found that the number of *Rnd2*-deleted cells was significantly lower at 21 dpi compared to control cells (Fig. 2E; t_7_=3.71, p=0.008). Importantly, when we performed the same analysis in wild-type C57Bl6/J mice (Fig. S4A), there was no difference between the two categories of cells, thus excluding the possibility that the observed effect is due to Cre toxicity. We further confirmed that the exacerbated cell death is specific to *Rnd2* suppression since the survival of *Rnd2*-knockout cells could be restored to levels which are not significantly different to control by co-delivery of a retroviral vector encoding Rnd2 expression construct (Fig. S4B, C). In contrast the loss of *Rnd2* knockout cells could not be restored by co-delivery of dominant negative (DN)-RhoA (Fig. S4B, C). Thus, suppression of *Rnd2* in adult-born DGNs impairs their survival.

### Rnd2 controls the positioning, the morphogenesis and the functional maturation of adult-born DGNs

We next examined the role of Rnd2 in the migration of adult-born DGNs. Previous studies have established that adult-born DGNs migrate into the GCL during the second week after birth (Esposito et al., 2005) and they contribute mostly to the inner third of the GCL and to a lesser extent to the mid third (Duan et al., 2007; Kempermann et al., 2003). To analyze the impact of *Rnd2* deletion on this developmental step, we determined the relative position of GFP and GFP/Cre retrovirus-labeled cells expressing DCX in the GCL at 7, 14 and 21 dpi (Fig. 3A). This relative position was examined as already described (Kerloch et al., 2018): the inner border of the GCL was defined as the baseline to measure cell migration and the perpendicular distance from this baseline to the center of each cell body was measured and normalized by the thickness of the GCL (Fig. 3B). At 21 dpi, we found that *Rnd2*-knockout neurons migrated a further distance in the GCL when compared to control neurons (Fig. 3C; t_8_=2.34, p=0.05), indicating that Rnd2 influences the positioning of newborn neurons in the GCL of the adult DG.

**Figure 3:**
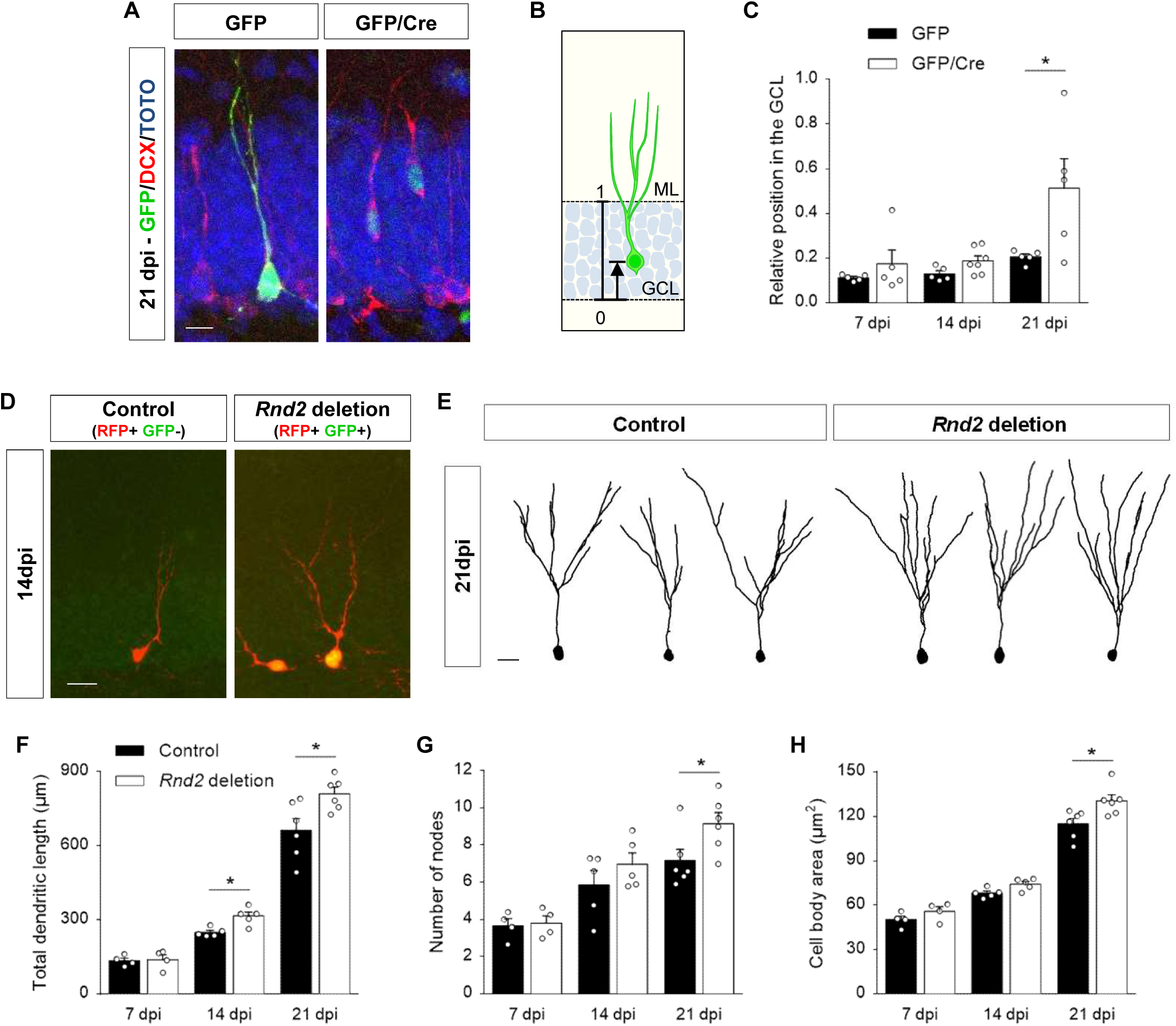
*Rnd2* deletion affects the positioning and the morphogenesis of adult-born DGNs. (**A**) Representative images of adult newborn neurons transduced with GFP or GFP/Cre retrovirus at 21 dpi in the DG. Doublecortin (DCX) identifies immature neurons and TOTO-3 labels nuclei. (**B**) Schematic depiction of the analysis of the relative positions of newborn neurons in the granular cell layer (GCL) of the DG. ML, molecular layer. (**C**) Relative positions of newborn neurons within the GCL in control group (GFP) and after *Rnd2* deletion (GFP/Cre). Mean ± s.e.m.; unpaired two-tailed Student’s t-test; *p < 0.05 (n = 5-7 mice, a minimum of 6 cells were analyzed per animal). (**D**) Representative images of control (RFP+ GFP-) and *Rnd2*-depleted (RFP+ GFP+) newborn neurons in the DG of *Rnd2*^*flox/flox*^ mice, 14 days after the injection of a mixture of GFP/Cre and RFP retroviruses. (**E**) Representative tracings of control and *Rnd2*-deleted newborn neurons in the adult DG at 21 dpi. (**F, G, H**) Quantification of the total dendritic length (F), the number of nodes (G) and the cell body area. Mean ± s.e.m.; paired two-tailed Student’s t-test; *p < 0.05, (n = 4-6 mice, a minimum of 4 cells were analyzed per animal). Scale bars represent 10 µm (A) and 20 µm (D, E).

While migrating, adult-born DGNs develop their dendritic arborization (Esposito et al., 2005). Three weeks after birth, their overall morphology resembles that of neurons at later time points (Zhao et al., 2006). To determine whether Rnd2 is also important for this process, we examined the morphology of newborn neurons in *Rnd2*^*flox/flox*^ brains injected with a mixture of GFP/Cre and RFP retroviruses. In the same animals, we were able to reconstruct the somatodendritic structure of control cells (cells expressing only RFP) and *Rnd2*-deficient cells (cells expressing GFP and RFP) (Fig. 3D). Interestingly, our results showed that *Rnd2*-deleted neurons have longer (Fig. 3E-F; t_4_=3.64, p=0.02 at 14 dpi; t_5_=3.36, p=0.02 at 21 dpi) and more branched dendrites compared to control neurons (Fig. 3E,G; t_5_=3.39, p=0.02 at 21 dpi). Furthermore, we found that cell bodies of *Rnd2*-deleted neurons were significantly larger than those of control neurons at 21 dpi (Fig. 3H; t_5_=3.17, p=0.02). We then analyzed the impact of *Rnd2* deletion on spine density and morphology at 21 and 28 dpi, a stage representing the peak of spine growth (Zhao et al., 2006). We observed that the total spine density of *Rnd2*-deficient newborn neurons was not statistically different compared to controls and no major difference in spine morphology was found between the two groups (Fig. S5). Altogether, these results show that, besides their correct distribution in the GCL, Rnd2 controls the morphogenesis of adult-born DGNs, including soma size and dendritic tree extent.

The aforementioned results then raised the following question: do *Rnd2*-deleted neurons die because of their mispositioning and/or aberrant morphology or because Rnd2 has a direct effect on the survival of adult newborn neurons? To discriminate between these two scenarios, we delivered a retrovirus expressing an inducible Cre (Mu et al., 2015) in order to delete *Rnd2* expression at a specific stage, in particular 3 weeks after birth, when new neurons have reached their final position and established their arborization (Fig. 4A). Using this approach, we found that *Rnd2* deletion led to a significant decrease in the survival of adult-born DGNs (Fig. 4B; t_12_=2.31, p=0.04), whereas their positioning and morphology were not perturbed (Fig. 4C-E), indicating that Rnd2 controls the survival of these cells independently of their position and morphology. Interestingly, while induction of *Rnd2* deletion at 28 dpi still led to a decrease of newborn neuron survival (Fig. S6; t_11_=2.75, p=0.02 for survival), *Rnd2* deletion at 56 dpi did not (Fig. 4F-J; t_12_=0.84, p=0.42 for survival). Therefore, these experiments suggest that Rnd2 is critical for adult hippocampal newborn neuron survival during a defined period of their development, at the immature stage.

**Figure 4:**
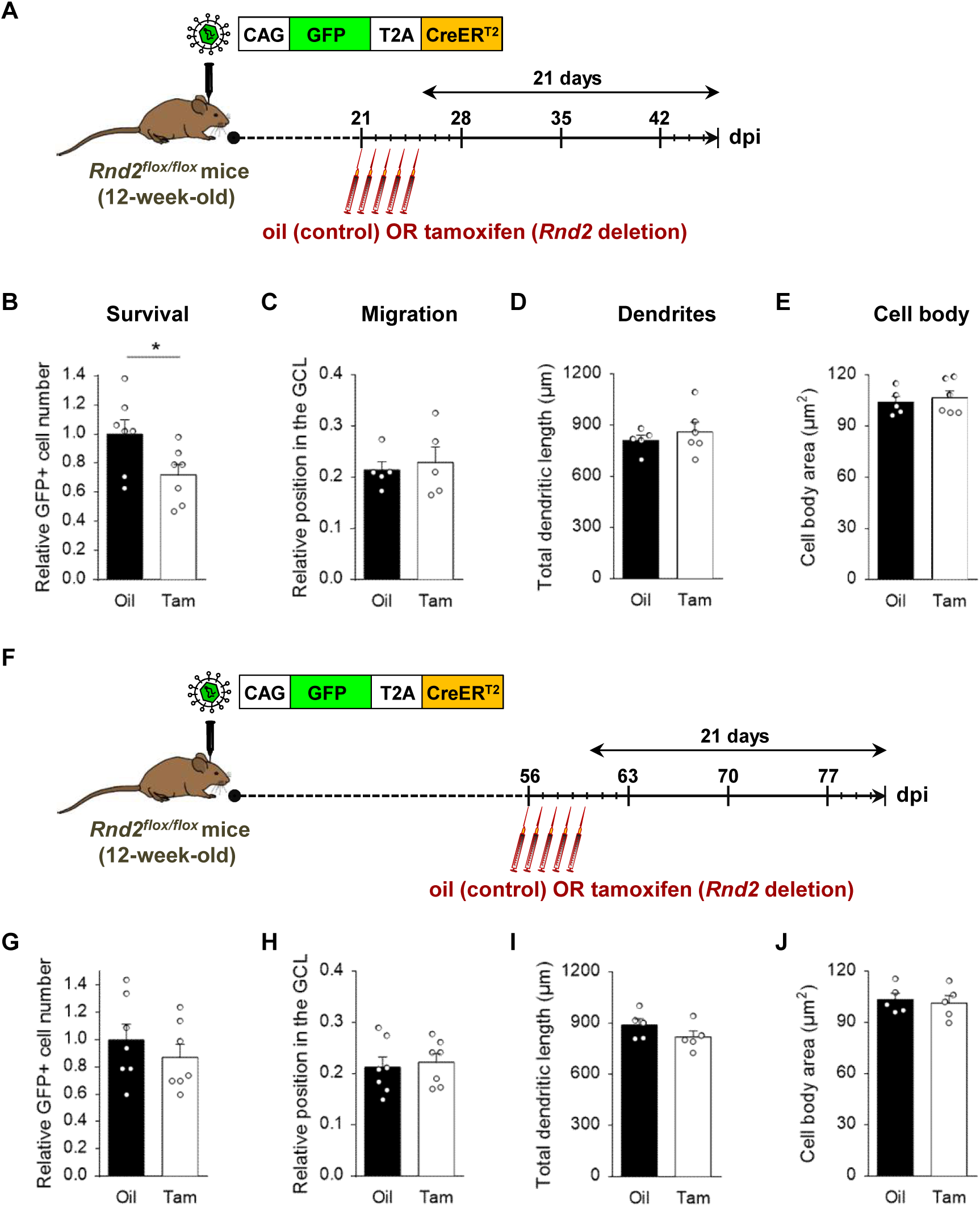
*Rnd2* deletion impacts directly the survival of adult-born DGNs during a specific period of their development. **(A, F**) Experimental designs. A retrovirus expressing GFP together with a conditionally active form of Cre recombinase, which is activated upon tamoxifen, was injected into the DG of adult *Rnd2*^*flox/flox*^ mice. Three (A) or eight weeks (F) after virus injection, tamoxifen (150 mg/kg, daily for 5 days), or oil in control group, was injected and animals were sacrificed 21 days after the last injection of tamoxifen. dpi, days post-injection. (**B-E, G-J**) The relative number of GFP+ cells, the relative position in the GCL and the morphology of transduced cells were quantified 21 days after the last injection of tamoxifen. Mean ± s.e.m., Unpaired two-tailed Student’s t-test; *p<0.05 (n=7 animals per group).

To further understand the impact of Cre-mediated *Rnd2* deletion in adult-born DGNs, we performed electrophysiological recordings of adult-born DGNs in acute slices from GFP or GFP/Cre virus-infused animals under whole-cell current-clamp configuration. Through this approach, we found that the membrane properties of 4 week-old adult-born DGNs (membrane resistance, capacitance and resting potential) were not affected by the suppression of *Rnd2* (Fig. S7A-D). In response to a depolarizing current step, we also examined the ability of new neurons to generate action potentials (APs), a hallmark of neuronal maturation. As compared to control, we found that the AP threshold was significantly increased in *Rnd2*-deleted neurons (Fig. 5A, B; −40.4 ± 1.4 mV versus −44.1 ± 0.5 mV; t_45_=2.98, p=0.005). Moreover, the AP amplitude was smaller in *Rnd2*-deficient newborn neurons (Fig. 5A, C; 82.1 ± 4.5 mV versus 96.4 ± 1.5 mV; t_45_=3.64, p=0.0007) and the maximal frequency of these neurons to fire APs tended to be reduced compared to control neurons (Fig. 5D; 35.1 ± 4.1 Hz versus 41.6 ± 1.5 Hz; t_45_=1.78, p=0.08), which altogether indicate that *Rnd2*-deleted newborn neurons have a decreased ability to fire APs. These results suggest that Rnd2 is critical for the proper development of intrinsic excitability of adult-born DGNs and consequently for their integration into the hippocampal circuitry.

**Figure 5:**
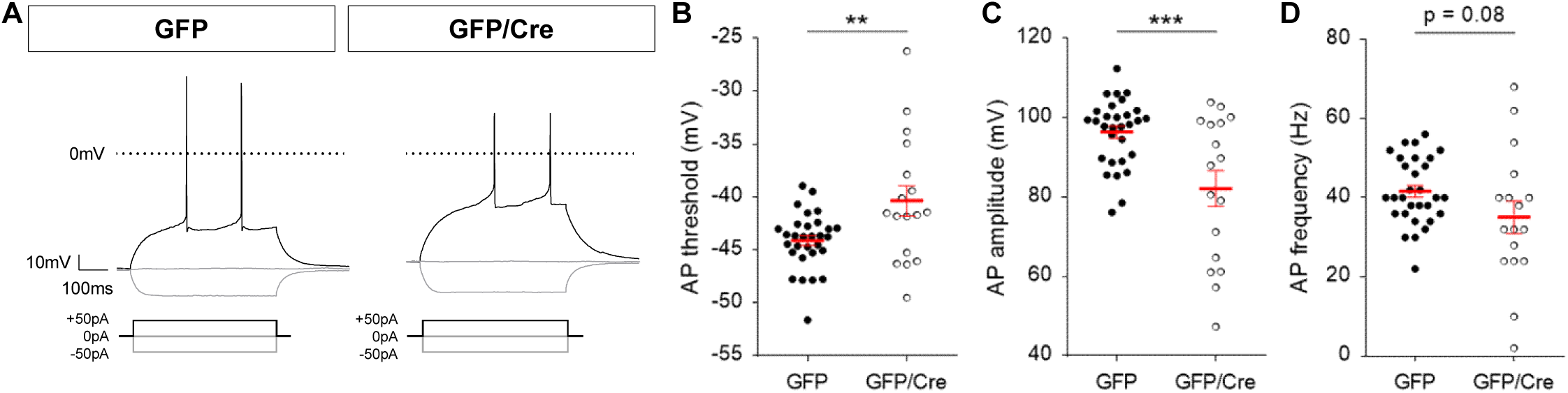
*Rnd2* deletion affects the intrinsic excitability of adult-born DGNs. (**A**) Representative spiking pattern of control (GFP) and *Rnd2*-deleted (GFP/Cre) newborn neurons in response to current injection (0 or ± 50 pA for 500 ms). (**B, C, D**) Quantification of the threshold of the first elicited AP (B), the maximum amplitude of this first AP (C) and the maximum AP frequency elicited by current injection (D). Mean ± s.e.m; unpaired two-tailed Student’s t-test; **p < 0.01, ***p < 0.001 (n = 30 neurons from 4 mice in GFP group and n = 17 neurons from 8 mice in GFP/Cre group).

### Rnd2 expression in adult-born DGNs controls anxiety-like behaviour

It has been reported that new neurons in the adult DG are of pivotal importance for several hippocampal-dependent functions (Toda et al., 2018) including spatial navigation (Dupret et al., 2008) as well as behavioural pattern separation (Clelland et al., 2009; Nakashiba et al., 2012; Tronel et al., 2010a), a process allowing to distinguish highly similar events, objects or contexts. In addition, AHN has been shown to be implicated in the regulation of affective states (Anacker and Hen, 2017), particularly in anxiety (Revest et al., 2009). Given our observation that Rnd2 is crucial for the survival of adult newborn neurons and their maturation within the DG, we studied whether the deletion of *Rnd2* specifically in these cells impairs hippocampal-dependent memory, anxiety and/or depression-like behaviours. In this goal, two batches of adult *Rnd2*^*flox/flox*^ mice were injected bilaterally into the DG with high-titer retroviruses expressing GFP or Cre/GFP and behaviourally tested at least four weeks after the injection (Fig. S8A).

To evaluate memory in these mice, we first studied spatial navigation according to a classical procedure in the Morris water maze (MWM) that allows to test reference memory (Fig. S8B). We found that spatial memory was intact following *Rnd2* deletion in adult-born DGNs (Fig. S8C). Moreover, no memory deficit was observed during the probe test, performed 3 days after the completion of the learning phase, indicating that memory retention and recall were intact (Fig. S8D). Next, to explore the impact of *Rnd2* deletion on behavioural pattern separation, mice were trained in a contextual fear conditioning paradigm to assess their ability to discriminate two similar contexts (Tronel et al., 2010a) (Fig. S8E). The results show that Cre-mediated deletion of *Rnd2* in adult-born DGNs did not impact contextual fear conditioning (Fig. S8F). In addition, no generalization was observed since both groups showed a lower freezing response in context B compared to context A, 24h or 5 weeks after conditioning (Fig. S8F, G). Thus, Cre-mediated deletion of *Rnd2* in adult-born DGNs does not affect behavioural pattern separation.

We next examined the impact of *Rnd2* suppression in adult-born DGNs on anxiety-like behaviour by measuring avoidance responses to potentially threatening situations, such as open and bright environments. Using the open field test, we found, 4 weeks after retrovirus injection, that *Rnd2*-deficient mice showed comparable locomotor activity compared to control mice (Fig. S9A). In this test, the time spent in the center was not significantly different between the two groups (Fig. S9B, t_25_=0.64, p=0.53). Following this test, mice were subjected to the emergence test using a dark cylinder placed into a bright open field (Fig. S9C-E). In this task, we observed no significant difference between groups although the latency to emerge from the reassuring cylinder tends to be longer for *Rnd2*-deficient mice compared to control mice (Fig. S9D, t_25_=1.73, p=0.10). In contrast, in the elevated plus maze (EPM), *Rnd2*-deficient mice (GFP/Cre) spent significantly less time in the open arms, which are more threatening areas, compared to control mice (GFP) (EPM1, Fig. 6A, t_24_=2.48, p=0.02). Interestingly, the reduction of time spent by GFP/Cre-infused mice in the open arms of the EPM remained significantly decreased compared with control mice as long as 10 weeks after virus injection, suggesting a long-lasting effect of the deletion (EPM2, Fig. 6A; t_25_=2.19, p=0.04). Importantly, this effect was not due to a modification of locomotor activity and/or exploration since the number of total entries was similar between the two groups (Fig. 6B). Consistent with these observations, *Rnd2*-deficient mice also exhibited a significant increase in anxiety-like behaviour when subject to a light/dark test. In this test, *Rnd2*-deleted mice escaped more quickly from the illuminated chamber (Fig. 6C, t_18_=2.80, p=0.01) and spent significantly less time in this chamber compared to control mice (Fig. 6D, t_18_=4.63, p<0.001).

**Figure 6:**
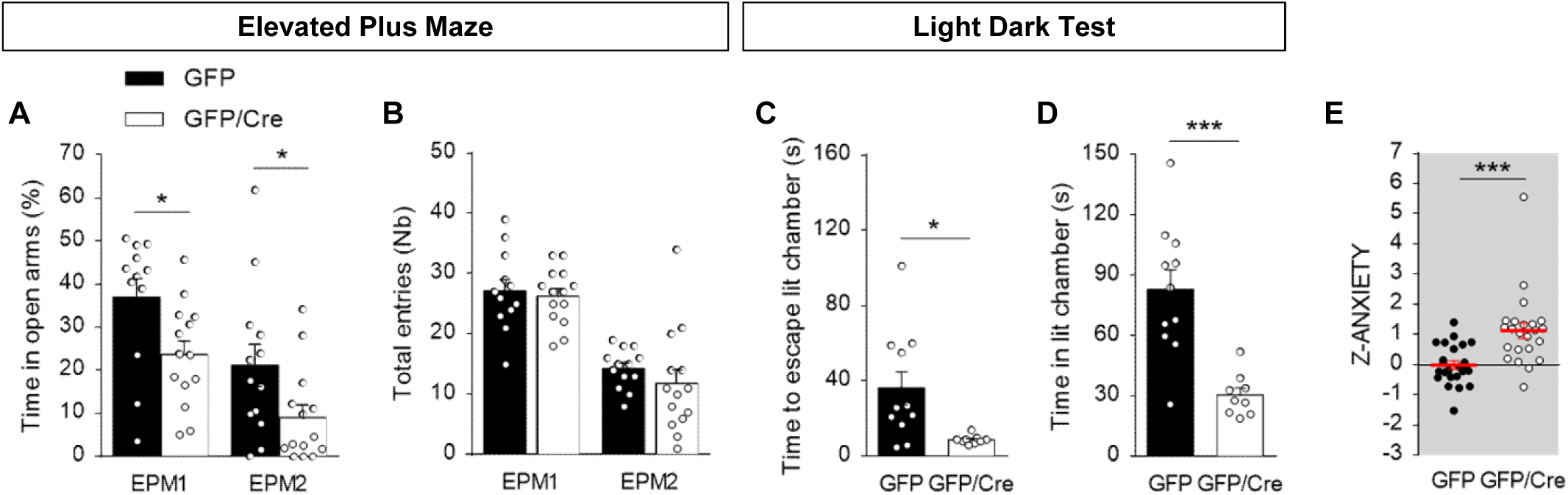
*Rnd2* deletion in adult-born DGNs increases anxiety-like behaviour. (**A**) Time spent in the open arms and (**B**) number of total entries in the elevated plus maze (EPM), 4 (EPM1) or 10 weeks (EPM2) after GFP or GFP/Cre retrovirus injection. Mean ± s.e.m., Unpaired two-tailed Student’s t-test; *p<0.05. (**C**) Time to escape the lit chamber in the light/dark test and (**D**) time spent in this chamber 5 weeks after retroviral injection. Mean ± s.e.m., Unpaired two-tailed Student’s t-test; *p<0.05, ***p<0.001. (**E**) z-anxiety score calculated by averaging z-score values of the different tests used to assess anxiety-like behaviour. An increased score value reflects increased anxiety. Mean ± s.e.m., Unpaired two-tailed Student’s t-test; ***p<0.001.

In a subsequent analysis, a z-normalization was applied (Guilloux et al., 2011). This methodology standardizes observations obtained at different times and from different cohorts, thus allowing their compilation. For each anxiety-like behaviour, a Z-score was calculated (see Methods). The directionality of scores was adjusted such that increased score values reflect increased anxiety. Then the z values obtained for each test were averaged to obtain a single z-anxiety score. As shown in Fig. 6E (t_45_=4.01, p<0.001), this score is significantly increased in mice injected with the GFP/Cre retrovirus compared to control mice further supporting that the suppression of *Rnd2* in adult-born DGNs increases anxiety-like behaviour.

In addition to studying anxiety behaviour, we also analyzed depression-like behaviour. Using a sucrose preference test, we found that the total liquid intake as well as the consumption of sucrose were similar between the two groups indicating that the deletion of *Rnd2* does not induce anhedonia-like behaviour (Fig. S9F-G). Next, we assessed immobility during exposure to inescapable stress using the forced-swim test (FST). In this test, we found that both the latency to immobility and the total time spent floating were not significantly modified by the suppression of *Rnd2*, despite a trend (Fig. S9H-I, t_18_=1.92, p=0.07 for immobility time). When Z-score normalization was performed (Fig. S9J), we found no significant difference between the two groups suggesting that the absence of *Rnd2* in newborn neurons of the adult DG does not impact depression-like behaviour.

Finally, the number and the position of adult-born DGNs targeted by our retroviral approach were quantified at the end of the behavioural sequences. In both batches of animals, we confirmed that the deletion of *Rnd2* increases the death of newborn neurons in the adult DG since fewer GFP+ cells were detected in mice injected with the GFP/Cre retrovirus compared to control mice injected with the GFP retrovirus (Fig. S9K-N), although similar titers of virus were infused (see Methods). In addition, transduced cells were located along the septo-temporal axis of the DG. Altogether, these experiments demonstrate that *Rnd2* deletion in adult-born DGNs via a retroviral approach impacts anxiety-like behaviour but does not affect memory and depression-like behavior.

### Rnd2 plays distinct functions in developmentally- and adult-born DGNs

Lastly, we asked whether the functions of Rnd2 described so far are specific to adult-born DGNs or, in other words, whether Rnd2 plays similar roles in developmentally-born DGNs. To address this, *Rnd2* was deleted using a retroviral approach identical to the one used in adult mice but in this case GFP or GFP/Cre retroviruses were injected at P0 when generation of DGNs reaches a peak in mice (Snyder, 2019). Importantly, *Rnd2* is prominently expressed at this stage in the DG (Fig. 7A). Since most defects were observed at 21 dpi in adult-born DGNs, a similar time point of analysis was used in this set of experiments. Moreover we only focused on the cellular processes that were affected in adult-born DGNs, i.e. survival, positioning and somatodendritic morphogenesis. First, to study the impact of the deletion on cell survival, we looked at caspase-3a expression. Although we were able to detect some caspase-3a+ cells in the DG at 21 dpi (Fig. 7B), we never detected any GFP cells expressing this apoptotic marker in both conditions (at least 100 GFP+ cells from 3-4 animals were analyzed in each group). We also performed this analysis at a later time point, 12 weeks post-injection (wpi), since it has been suggested that developmentally-born DGNs, in contrast to adult-born DGNs, do not go through a period of cell death during their immature stages but instead die after reaching maturity, between 2 and 6 months of age (Cahill et al., 2017). However, even at this stage, no control or *Rnd2*-deleted GFP+ cells were found to express caspase-3a (data not shown). Based upon our observations, the survival of P0-born DGNs is not significantly affected by the absence of *Rnd2*, a finding which contrasts the effect of *Rnd2* deletion in adult-born DGNs. We looked further in these studies and found that the final position of this population of neurons was not impacted by *Rnd2* suppression (Fig. 7C, D). For morphological analysis, RFP and GFP/Cre retroviruses were co-injected at P0 similarly to our experiments in adult mice. Through this analysis, we found that the size of the cell body was not significantly different between the two groups at 21 dpi and 12 wpi (Fig. 7E, F). However the total dendritic length (Fig. 7E, G; t_3_=4.06, p=0.03 at 21 dpi; t_3_=6.26, p=0.008 at 12 wpi) as well as the number of nodes (Fig. 7E, H; t_3_=7.33, p=0.005 at 21 dpi; t_3_=3.57, p=0.04 at 12 wpi) were both significantly increased in *Rnd2*-depleted neurons compared to control neurons, when measured at both 21 dpi and 12 wpi. Finally, we assessed whether *Rnd2* deletion in neonatally-born DGNs also affect anxiety-like behaviour. In this goal, P0 *Rnd2*^*flox/flox*^ pups were injected bilaterally with Cre/GFP or GFP retrovirus and tested, once they reached adulthood, using the same behavioural tasks as those described for our aforementioned studies of adult-born DGNs (Fig. S10A). Interestingly, the suppression of *Rnd2* in neonatally-born DGNs has no impact on anxiety-related behaviour (Fig. 7I-M, S10), suggesting that Rnd2 has specific and unique functions in DGNs generated during adulthood. To conclude, in neonatally-born DGNs, Rnd2 seems to be crucial only for dendrite morphogenesis, indicating that granule neurons in the DG exhibit a differential dependency to Rnd2 according to the age and the environment of the hippocampal neurogenic niche.

**Figure 7:**
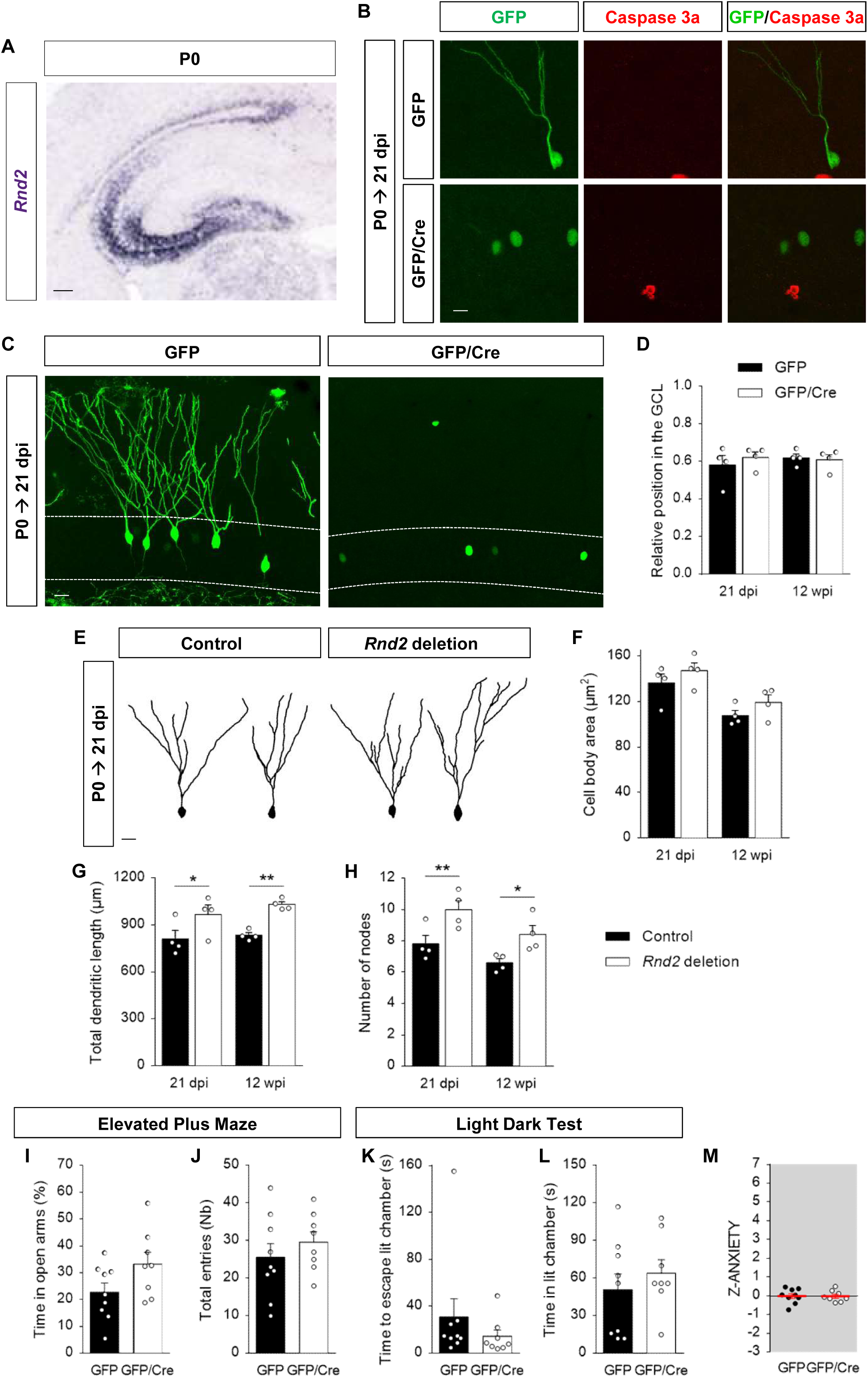
*Rnd2* suppression in neonatally-born DGNs only affects their dendritic development. (**A**) Distribution of *Rnd2* transcripts in the mouse hippocampus at P0. (**B**) Immunostaining for GFP and caspase-3a in the DG of *Rnd2*^*flox/flox*^ pups, 21 days after GFP or GFP/Cre retrovirus injection at P0. (**C**) Representative images and (**D**) quantification of the relative position of GFP+ neurons within the GCL, 21 days after GFP or GFP/Cre retrovirus injection at P0. Mean ± s.e.m. (n=4 mice). (**E**) Representative tracings of control and *Rnd2*-deleted neonatally-born DGNs at 21 dpi. (**F, G, H**) Quantification of the cell body area (**F**), the total dendritic length (**G**) and the number of nodes (**H**) 21 days after P0 injection. Mean ± s.e.m.; paired two-tailed Student’s t-test; *p < 0.05, **p < 0.01 (n=4 mice, a minimum of 3 cells were analyzed per animal). (**I**) Time spent in the open arms and (**J**) number of total entries in the elevated plus maze. (**K**) Time to escape the light room in the light/dark test and (**L**) time spent in this room. (**M**) z-anxiety score calculated by averaging z-score values of the different tests used to assess anxiety-like behavior. Mean ± s.e.m., GFP n=9 and GFP/Cre n=8 mice. Scale bars represent 100 µm (A), 10 µm (B) and 20 µm (C, E).

## DISCUSSION

Despite the central role of Rho GTPases in neuronal development (Govek et al., 2005), their function in adult neurogenesis has been poorly investigated (Vadodaria and Jessberger, 2013) and the role in particular of the Rnd subfamily is totally unknown. In this study, by using an approach of loss of function, we provide evidence that Rnd2 is essential for AHN. At the cellular level, we demonstrate that Rnd2 is cell-intrinsically required for the survival and maturation of adult-born DGNs. At the organism level, we show that the expression of Rnd2 in adult-born DGNs is critical for the control of anxiety behaviour. Importantly, we also reveal that Rnd2 differentially regulates adult and developmental neurogenesis in the DG since its deletion in DGNs generated at birth only affects their dendritic arbor. Our work thus highlights new important functions for this atypical Rho GTPase *in vivo*.

### Rnd2 has unique functions in the development of adult-born DGNs

So far, our knowledge on Rnd2 functions in neurons *in vivo* is based on studies performed in the context of cerebral cortex development (Heng et al., 2008; Pacary et al., 2011). During corticogenesis, Rnd2 has been shown to control neuronal migration (Heng et al., 2008; Pacary et al., 2011) and neurite outgrowth (Heng et al., 2008). The finding that Rnd2 is a key player in newborn neuron positioning and morphology in the adult DG was thus not surprising. In contrast, the finding that Rnd2 is crucial for the survival of adult-born DGNs was more unexpected because Rnd2 has never been described, to our knowledge, as a regulator of cell survival in any cell types. This function of Rnd2 in adult newborn neurons seems thus unique. In contrast to Rnd2, Rnd3 has been shown to modulate cell survival and apoptosis (Riou et al., 2010). Moreover, Rnd3 is involved in neural progenitor proliferation (Pacary et al., 2013), neuronal migration (Azzarelli et al., 2014; Pacary et al., 2011) and polarization (Peris et al., 2012) whereas Rnd1 regulates late stages of neuronal development such as dendrite and spine formation (Ishikawa et al., 2003, 2006). Therefore, it will be interesting in future experiments to determine whether the two other Rnds also contribute to AHN and whether they have similar functions. However, in view of their distinct expression pattern in the adult DG (Fig. S1), we can already speculate that Rnd1 and Rnd3 might mediate different functions than Rnd2 in this process. Moreover, Rnd1 and Rnd3 have been shown, in different cell types including neurons, to be predominantly expressed at the plasma membrane whereas Rnd2 is mainly located in endosomes (Pacary et al., 2011; Roberts et al., 2008). Such differences in subcellular localization further supports our hypothesis that Rnd2 might dictate different functions than Rnd1 and Rnd3 in adult-born DGNs.

Our data obtained with the retrovirus expressing an inducible Cre (Fig. 4, S6) indicate that Rnd2 controls adult newborn neuron survival independently of its role in their positioning and morphology. Interestingly, when *DISC1* (Disrupted-In-Schizophrenia 1) was downregulated in adult-born DGNs using a similar retroviral approach, Duan et al. observed an ectopic migration of these neurons in the molecular layer, an increase of cell body size and dendrite length, similarly to the phenotypes observed after *Rnd2* deletion, but no increased cell death (Duan et al., 2007). This study thus suggested that an abnormal morphological development and an aberrant positioning of adult-born DGNs do not result in the death of these cells further reinforcing that the control of newborn survival is independent of the regulation of their position and morphology.

Indirectly, this set of experiments with the inducible Cre also suggests that Rnd2 regulates newborn neuron survival and their maturation through distinct mechanisms. The signaling pathways mediating Rnd2 action in general are poorly understood (Azzarelli et al., 2015). In cortical neurons, during embryogenesis, Rnd2 promotes migration independently of the actin cytoskeleton but partially through the inhibition of RhoA (Pacary et al., 2011) and pharmacological inhibition of RhoA signaling was shown to enhance the survival of adult-born DGNs (Christie et al., 2013). However, we found that the excess death of *Rnd2*-deleted adult-born DGNs was not counteracted by co-delivery of DN-RhoA, suggesting that the action of Rnd2 on adult newborn survival does not involve RhoA inhibition (Fig. S4B, C). Other candidate interactors might be relevant to the functions of Rnd2 in adult-born DGNs. For example, semaphorin receptors, known as Plexins, have been implicated in AHN (Duan et al., 2014; Jongbloets et al., 2017; Mata et al., 2018; Zhao et al., 2018) and have been shown to be bound and regulated by Rnd2 in neurons and other cell types (Azzarelli et al., 2014; McColl et al., 2016; Uesugi et al., 2009). Nevertheless, whether Plexins mediate some aspects of Rnd2 action in adult newborn neurons remains unknown.

As mentioned previously, Rnd2 has been shown to be expressed in endosomes (Pacary et al., 2011; Roberts et al., 2008; Tanaka et al., 2002) and to interact with molecules involved in the formation and trafficking of endocytic vesicles (Kamioka et al., 2004; Tanaka et al., 2002). In the adult DG, our immunostainings (Fig. 1F-I) suggest that Rnd2 might be also localized to this subcellular compartment. This raises the possibility that Rnd2 activity in adult newborn neurons may involve endocytosis and, in particular, Rnd2 may regulate the trafficking of membrane-associated molecules such as receptors or adhesion molecules that control survival, migration and/or morphological development. Among these molecules, the NMDA receptor could be a candidate. Indeed, like Rnd2, NMDA receptors are critical for the survival of immature neurons at the time of their integration into the pre-existing neuronal network (Tashiro et al., 2006a).

### Rnd2 differentially regulates neurogenesis in the adult and developing DG

Adult neurogenesis is often viewed as a continuation of developmental neurogenesis. However, although adult neurogenesis recapitulates the entire process of neuronal development that occurs during embryonic or early postnatal stages, fundamental differences are evident between development and adulthood (Gotz et al., 2016; Urban and Guillemot, 2014). For example, in the developing brain, nascent neurons must cope with a continuously changing environment, while their counterparts in adult brain are surrounded by a relatively stable niche. More specifically in the DG, adult-born DGNs and developmentally-born DGNs exhibit distinct morphological features once mature (Cole et al., 2020; Kerloch et al., 2018), undergo distinct survival dynamics (Cahill et al., 2017), have differential ability to reshape in response to a seizure or learning experience (Kron et al., 2010; Lemaire et al., 2012; Tronel et al., 2010b), are recruited during different learning tasks (Tronel et al., 2015) and are required for different functions (Nakashiba et al., 2012; Wei et al., 2011). However, despite such clear differences, the intrinsic molecular mechanisms underlying the distinct regulation of developmental and adult neurogenesis remain largely unknown. Here, by demonstrating that Rnd2 regulates differently the development of DGNs generated in the early postnatal brain versus young adult brain, we have identified a key factor for this differential regulation. This further argues that adult neurogenesis is not a simple continuation of earlier processes from development. How Rnd2 plays such distinct functions in developmentally-born and adult-born DGNs remains to be elucidated. The absence of specific interactors/effectors of Rnd2 during development might be part of the answer.

### Rnd2 in adult-born DGNs: a new target for anxiety-like disorders

We show here that *Rnd2* deletion in adult-born DGNs impacts anxiety-like behaviour while depression-like behaviour is not affected. These results reproduce what we and others have obtained following depletion of new DGNs in the adult brain. Indeed, transgenic mice (Nestin-rtTA/TRE-Bax mice), in which AHN is specifically impaired by over-expressing the pro-apototic protein Bax in Nestin+ cells of the DG, exhibit a striking increase in anxiety-related behaviour whereas depression like-behaviour remains unchanged (Revest et al., 2009). Similarly, *TrkB* deletion in adult hippocampal progenitors decreases newborn neuron survival in the adult DG and promotes an anxiety-like state without affecting depression (Bergami et al., 2008). Consistent with these findings, it is becoming clear that AHN cannot be considered as an etiological factor of depression, but rather as a mediator of the effects of antidepressants or the effect of stress on depression-like behavior (Anacker et al., 2018; Eliwa et al., 2017; Miller and Hen, 2015; Santarelli et al., 2003).

Moreover, in our study, reference memory and behavioral pattern separation are not altered. These results were unexpected since we showed that these processes are impaired in the Nestin-rtTA/TRE-Bax mice (Dupret et al., 2008; Tronel et al., 2010a). Several factors may explain these distinct results, including differences in the experimental design of behavior assays or the method used to target adult newborn neurons. For example, the retroviral strategy only targets a population of new neurons born at a specific time whereas the previous approach, using doxycycline for ablation, targets a broader population. So memory processes might be affected only when a large population of adult newborn DGNs is suppressed. In accordance with this assumption, it was shown that an extensive lesion of the hippocampus is required to observe memory deficits (Moser et al., 1995). Consequently, anxiety-like behavior may be more sensitive to the loss of new neurons compared to memory processes, hence the behavior of *Rnd2-*deficient mice in the present study. An alternative explanation of our phenotype is that Rnd2 might play a more prominent role in adult-born DGNs located in the temporal DG. This hypothesis is based on several lines of evidence. Firstly, memory function depends on the septal hippocampus whereas anxiety is related to its temporal part (Bannerman et al., 2004; Fanselow and Dong, 2010). Secondly, adult-born DGNs in the temporal DG seem to be especially important in stress-induced regulation of anxiety (Anacker et al., 2018). Thirdly, *Rnd2* expression is particularly enriched in the temporal DG (Fig. 1E and http://hipposeq.janelia.org).

While our results indicate that altered levels of *Rnd2* in adult-born DGNs results in increased anxiety-like behavior, it is still unclear whether this behavioral effect is due to the death of newborn neurons, their mispositioning, their abnormal morphology, or all of these traits. Nevertheless, since *Rnd2* suppression in neonatally-born DGNs has no effect on anxiety-like behavior whereas their dendrites are affected, these results might be interpreted to suggest that the increased anxiety-like behavior following *Rnd2* loss in adult-born DGNs is not due to the abnormal dendritic arbor of these cells.

Overall, our findings demonstrate that Rnd2 is an essential molecular player for the proper development of adult-born DGNs and, because of this function, serves a critical role in the regulation of anxiety-like behavior. These data validate not only the neurogenesis hypothesis of anxiety but also identify Rnd2 as a molecular link between AHN and anxiety. Targeted manipulation of Rnd2 in adult-born hippocampal neurons might be thus a potential novel strategy for anxiety disorders.

## MATERIALS AND METHODS

### Animals and genotyping

C57Bl6/J (Janvier) and *Rnd2*^*flox/flox*^ mice were housed, bred, and treated according to the European directive 2010/63/EU and French laws on animal experimentation. All procedures involving animal experimentation and experimental protocols were approved by the Animal Care Committee of Bordeaux (CEEA50) and the French Ministry of Higher Education, Research and Innovation (authorizations n°04997.02 and APAFIS*#*12546).

*Rnd2* conditional mutant allele is described in Figure S2A. *Rnd2* was targeted by homologous recombination to generate the *Rnd2*^*flox*^ allele. This allele contains one loxP site between the exon 1 and 2 and a second loxP site after the last exon. A neomycin (Neo) selection gene flanked by flippase recognition target (FRT) sites was inserted in 3’ of *Rnd2*. Following transmission of the mutation to the germline, the Neo gene was excised, giving rise to the *Rnd2*^*flox*^ allele. Upon Cre recombination, exons 2 to 5 of *Rnd2* are deleted.

Genotyping of *Rnd2* wild-type allele was performed with the following primers: forward, 5’-CAGGGCACTTCTGATACAAAGC -3’ and reverse 5’-TCTCACCCACCCCTGGCTGAT - 3’. Genotyping of *Rnd2* conditional mutant allele was performed with the same forward primer as for the wild-type allele (WT) and the following reverse primer 5’-GTTTGTCCTCAACCGCGAGCTG -3’. Mice were genotyped from genomic DNA purified from tail biopsies by PCR using these primers according to the following protocol. Tails were incubated overnight at 56°C in Proteinase K (PK) buffer (100 mM Tris-HCl pH8, 5 mM EDTA, 0.2 % SDS, 200 mM NaCl, 0.2 mg/mL PK). After a centrifugation at 13200 rpm for 10 min, the supernatants were purified by vacuum on silica columns, according to the manufacturer’s protocol (Macherey-Nagel) and on Zephyr automatic station (Perkin-Elmer). PCR assay was carried out on a Bio-Rad C1000 thermal cycler, in a 20 µL volume, using GoTaq G2 Hot Start Green Master Mix (Promega), and 0.2 µM of common forward primer, 0.2 µM of flox Reverse primer, and 0.4 µM of WT reverse primer. PCR conditions were as follows: 1 cycle, 5 min at 95°C; 37 cycles, 30 sec at 95°C, 30 sec at 60°C, 45 sec at 72°C; 1 cycle, 5 min at 72°C. PCR products were analyzed on a Labchip GX microfluidic electrophoresis system (Perkin-Elmer) using the DNA5k kit.

### Constructs

The murine Moloney leukemia virus-based retroviral vectors CAG–GFP (Zhao et al., 2006), CAG-GFP/Cre (Tashiro et al., 2006a), CAG-RFP (Laplagne et al., 2006) and CAG-GFP-T2A-CreER^T2^ (Mu et al., 2015) were kind gifts from Dr Fred Gage and the retroviral CAG-IRES-DsRed vector (Heinrich et al., 2011) from Dr Benedikt Berninger. Rnd2 and DNRhoA were cloned by PCR using pNeuroD1-Rnd2 (Heng et al., 2008) and pRK5-myc-RhoA-T19N (Addgene 12963) as templates respectively and then inserted into the SfiI/PmeI sites of the CAG-Neurog2-IRES-DsRed retroviral vector (Heinrich et al., 2011) to generate CAG-Rnd2-IRES-DsRed and CAG-DNRhoA-IRES-DsRed.

### Retrovirus production and injection into the mouse DG

High-titers of retroviruses were prepared as previously published (Kerloch et al., 2018) with a human 293-derived retroviral packaging cell line (293GPG) kindly provided by Dr Dieter Chichung Lie. For experiments in adults, the retroviral solution (10^9^-10^10^ TU/ml) was injected into the DG of 12-week-old male *Rnd2*^*flox/flox*^ mice. Mice were anaesthetized by intraperitoneal injection of ketamine (120 mg/kg; Imalgene® 1000, Merial)-xylazine (16 mg/kg; Rompun®, Bayer HealthCare) mixture and eye ointment was applied to prevent eyes from over-drying. A cranial subcutaneous injection of 0.1 μl of lidocaine (20 mg/ml; Lurocaïne®, Vetoquinol) and a dorsal subcutaneous injection of 0.1 μl of meloxicam (0.5 mg/ml; Metacam®, Boehringer Ingelheim) were performed before settling the mouse into the stereotaxic frame. Betadine was applied, then the skin was cut and a pulled microcapillary glass tube (1-5 μL, Sigma) was placed above bregma. Coordinates of the injection site from bregma were: anteroposterior: −2 mm, mediolateral: −1.8 mm, dorsoventral: −2.2 mm. A small hole was made on the skull using an electric drill, the microcapillary was loaded with the retroviral solution and introduced within the hole to reach the DG and stayed in place for one minute. Then 1.5 μl of retrovirus was injected at the rate of 0.5 μl every two minutes and the microcapillary was left again in the same position for two minutes after the end of infusion before being removed. Then the skin was stitched with absorbable sutures and the mouse was placed in a recovery chamber (37°C) until it woke up. For injections in pups, postnatal day 0 (P0) mice were anesthetized by hypothermia and 1 µl of retroviral solution (10^9^ TU/ml) was injected through the skin and skull into the lateral ventricle using pulled borosilicate needles and a Femtojet microinjector. To target the lateral ventricle, a virtual line connecting the right eye with the lambda was used and the needle was injected slightly caudal of the midpoint of this line as previously described (Boutin et al., 2008). Injected pups were placed in a recovery chamber at 37°C for several minutes and then returned to their mother.

### Tamoxifen administration

For activation of the CreER^T2^ recombinase, animals were administered intraperitoneally with 150 mg/kg tamoxifen (Sigma, diluted in corn oil) for 5 consecutive days. Control animals were injected with the same volume of corn oil for 5 consecutive days.

### Tissue processing

Mice were deeply anesthetized with an intraperitoneal injection of pentobarbital (100 mg/kg; Pentobarbital® sodique, Ceva) and transcardially perfused with phosphate buffer saline (PBS, 0.1M, pH=7.3) and then with 4% paraformaldehyde (PFA) in PBS. Brains were dissected out of the skull, post-fixed in 4% PFA and cut coronally (40 µm) with a vibratome (Leica).

### Immunostaining

Rnd2 immunostaining was performed on sections obtained from freshly perfused brains and post-fixed 2 h with 4% PFA. After 3 washings in PBS, sections were treated with PBS - NH_4_Cl 50 mM for 15 min and blocked with a solution containing PBS - 0,05% saponin - 0,2% bovine serum albumin (BSA) for 15 min. They were then incubated overnight at 4°C with a goat anti-Rnd2 antibody (1/50, Santa Cruz, sc-1945) diluted in the blocking buffer. The following day, sections were washed three times in PBS – 0.05% saponin and incubated for 2 h with a donkey anti-goat Alexa Fluor® 488 (1/1000, Invitrogen A11055) diluted in PBS – 0.05% saponin – 0.2% BSA. For Rnd2/DCX immunostaining, a rabbit anti-doublecortin (1/2000, Sigma, D9818) was used.

For other immunostainings, each brain, after fixation and post-fixation with 4% PFA, was sectioned serially (10 series of 40 µm sections). For each staining, one in 10 free-floating sections (so one series) were used. After washing in PBS, sections were treated with PBS - 0.3% Triton X100 - 3% normal serum for 45 min. They were then incubated overnight at 4°C with the following primary antibodies diluted in PBS - 0.3% Triton X100 - 1% normal serum: rabbit anti-cleaved caspase 3 (1/400, Cell Signaling, 9661), rabbit anti-doublecortin (1/2000, Sigma, D9818), chicken anti-GFP (1/1000; Abcam, ab13970), rabbit anti-Ki67 (1/1000; NovoCastra, NCL-Ki67-P), rabbit anti-DsRed (1/500, Clontech, 632496). Sections were then incubated for 2 h at room temperature with appropriate fluorescent secondary antibodies diluted in PBS – 1% normal serum. TOTO-3 iodide (1/2000, Invitrogen) was added to the secondary antibody solution to label cell nuclei (Invitrogen). Images were acquired with a confocal microscope (Leica SP5 or SP8).

### Morphometric analysis

Dendrites and cell body of fluorescent dentate neurons were traced using a 100X objective and a semiautomatic neuron tracing system (Neuron Tracing from Neurolucida® software; MicroBrigthField). Measurements of node numbers, dendritic length and cell body area were performed with Neurolucida software.

For spine analysis, confocal stacks of images were obtained with a SP5 confocal microscope (63X oil-immersion objective; XY dimensions: 41.0 µm; z-axis interval: 0.13 µm). The dendritic length of each segment was measured on Z projections, and the number of dendritic spines was counted using NeuronStudio software (Rodriguez et al., 2008). Before spine analysis, images were deconvoluted using AutoQuantX3 software (Media Cybernetics). A minimum of 30 dendritic segments per experimental group and time point were examined for spine analysis.

### RNA in situ hybridization

Brains, freshly perfused and post-fixed overnight with 4% PFA, were cut with a vibratome (40 µm). Nonradioactive RNA in situ hybridizations on floating brain sections were then immediately performed with digoxigenin-labelled riboprobes as previously described (Azzarelli et al., 2014). The riboprobes used to visualize the expression of *Rnd2* (Heng et al., 2008) and *Rnd3* (Pacary et al., 2011) were previously described and the antisense RNA probe for *Rnd1* was prepared from IMAGE:3416797, GenBank accession number BE852181.

### Microdissection of the DG

Coronal sections (50 µm) were cut from frozen brains in isopentane using a cryostat (CM3050 S Leica) at −20°C and mounted on polyethylene-naphthalate membrane 1mm glass slides (P.A.L.M. Microlaser Technologies AG) that were pretreated to inactivate RNases. Sections were then immediately fixed for 30 sec with 95% ethanol and incubated with 75% ethanol for 30 sec and with 50% ethanol for 30 sec. Sections were stained with 1% cresyl violet in 50% ethanol for 30 sec and dehydrated in 50%, 75% and 95% ethanol for 30 sec each, and finally 2 incubations in 100% ethanol for 30 sec were performed. Laser Pressure Catapulting (LPC) microdissection of the DG (SGZ and GCL) (Fig. S1A) was performed using a PALM MicroBeam microdissection system version 4.6 equipped with the P.A.L.M. RoboSoftware (P.A.L.M. Microlaser Technologies AG). Laser power and duration were adjusted to optimize capture efficiency. Microdissection was performed at 5X magnification. Microdissected tissues were collected in adhesives caps and resuspended in 250 µl guanidine isothiocyanate-containing buffer (BL buffer from ReliaPrep™ RNA Cell Miniprep System, Promega) with 10 µl 1-Thioglycerol and stored at −80°C until RNA extraction was done. For the analysis of *Rnd* expression at different time points (Fig. 1D and S1B), the entire DG of each animal was microdissected. For the rostro-caudal analysis of *Rnd* expression (Fig. 1E and S1D), DG in the right hemisphere from 10 following coronal sections of 50 µm were pooled for each coordinate. In both cases, we began to microdissect the DG from the section at the anteroposterior coordinate −1.34 mm from the Bregma (when the two blades of the DG were visible).

Total RNAs were extracted from microdissected tissues using the ReliaPrep™ RNA Cell Miniprep System (Promega) according to the manufacturer’s protocol. The integrity of the RNA was checked by capillary electrophoresis using the RNA 6000 Pico Labchip kit and the Bioanalyser 2100 (Agilent Technologies), and quantity was estimated using a Nanodrop 1000 (ThermoScientific). The RNA integrity numbers (RIN) were between 8,2 to 10.

### Microdissection of GFP+ cells

For this analysis, *Rnd2*^*flox/flox*^ mice were injected bilaterally with 2 µl of high-titer retroviral solution. Brains were dissected out of the skulls in PBS and incubated in 30% sucrose / PBS for 3 days at 4°C under agitation. Brain were then frozen with isopentane and coronal sections (10 µm) were done using a cryostat and treated for microdissection as previously described but with some modifications. Sections were dehydrated in a series of pre-cooled ethanol baths (40 sec in 95% and twice 40 sec in 100%) and air-dried. Microdissection was performed at 63X magnification. Microdissected cells were collected in adhesives caps, resuspended in PK buffer (PK buffer from RNeasy FFPE Kit, Qiagen) and stored at −80°C until extraction was done. Total RNAs were extracted using the RNeasy® FFPE Kit (Qiagen,) according to the manufacturer’s protocol. The RNA integrity numbers (RIN) were above 7/8.

### Quantitative real-time PCR

RNA was processed and analyzed following an adaptation of published methods (Bustin et al., 2009). cDNA was synthesized from total RNA by using qSriptTM cDNA SuperMix (Quanta Biosciences). qPCR was performed using a LightCycler® 480 Real-Time PCR System (Roche). qPCR reactions were done in duplicate for each sample, using transcript-specific primers (Table 1), cDNA and LightCycler 480 SYBR Green I Master mix (Roche) in a final volume of 10 μl. PCR data were exported and analyzed in the GEASE software (Gene Expression Analysis Software Environment) developed in the Neurocentre Magendie (https://bioinfo.neurocentre-magendie.fr/outils/GEASE/). For the determination of the reference genes, the GeNorm method was used (Bustin et al., 2009). Relative expression analysis was corrected for PCR efficiency and normalized against two reference genes. The relative level of expression was calculated using the comparative (2-ΔΔCT) method (Livak and Schmittgen, 2001).

**Table 1:**
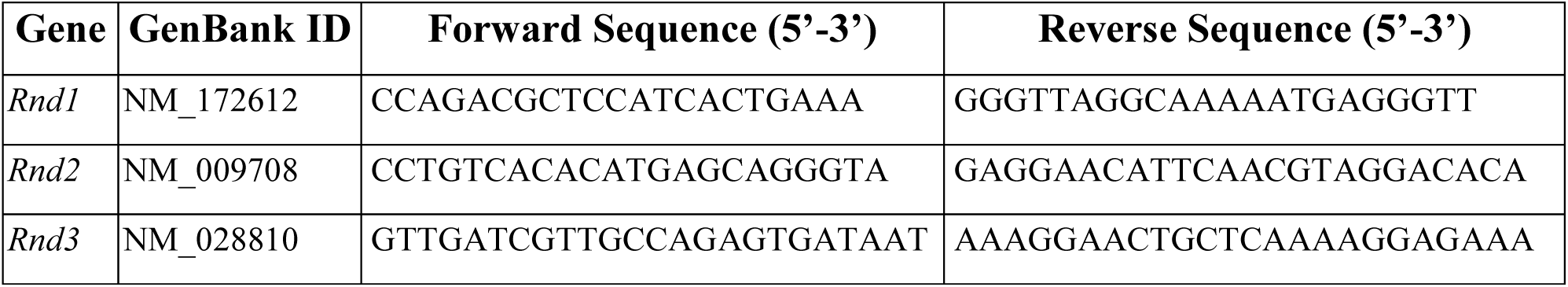
Sequences of primers used for qPCR.

### *Ex vivo* electrophysiology

Recombinant retroviruses encoding for GFP or GFP/Cre were bilaterally injected into the DG of two respective groups of 12-week-old *Rnd2*^*flox/flox*^ male mice. Four weeks after retroviral injection, animals were sacrificed by dislocation. The brains were quickly removed and immerged in ice-cold oxygenated cutting solution containing in mM: 180 Sucrose, 26 NaHCO_3_, 12 MgCl_2_, 11 Glucose, 2.5 KCl, 1.25 NaH_2_PO_4_, 0.2 CaCl_2_, oxygenated with 95% O_2_/5% CO_2_ ∼300 mOsm. Sagittal hippocampal slices (300 µm thick) were obtained using a vibratome (VT1200S, Leica) and transferred for 30 min into a 34°C bath of oxygenated ACSF containing in mM: 123 NaCl, 26 NaHCO_3_, 11 Glucose, 2.5 KCl, 2.5 CaCl_2_, 1.3 MgCl_2_, 1.25 NaH_2_PO_4_ ∼305 mOsm. After a minimum of 60 min recovery at RT (22-25°C), slices were transferred to a recording chamber in ACSF at 32°C. Recordings were performed using a Multiclamp 700B amplifier (Molecular devices) in fluorescent granule neurons clamped with glass pipettes (3-5 MΩ) filled with an internal solution containing in mM: 135 K-Gluconate, 10 KCl, 10 HEPES, 1 EGTA, 2 MgCl_2_, 0.3 CaCl_2_, 7 Phosphocreatin, 3 Mg-ATP, 0.3 Na-GTP; biocytin 0.4%; pH = 7.2; 290 mOsm. These cells were identified by their GFP expression, their morphology and soma location in the SGZ/GCL using a contrast microscope (Axio Examiner.A1, Zeiss) equipped with a fluorescent light (Colibri controller, Zeiss). Neuronal excitability was measured using 500 ms steps of current injections from −150 to 450 pA with increasing steps of 10 pA. Immunostaining of both GFP and biocytin was performed to confirm the identity of the recorded cells.

### Behaviour

For experiments with adults, 12-week-old male *Rnd2*^*flox/flox*^ mice were bilaterally injected with 1.5µl of GFP or GFP/Cre retroviruses on each side. Two batches of animals were used. The number of animals in each group was as follows: Batch 1 (GFP, 13 mice; GFP/Cre, 14 mice), Batch 2 (GFP, 11 mice; GFP/Cre, 9 mice). For the first batch, the titers of GFP and GFP/Cre retroviruses were 3-4.10^9^ TU/ml and for the second batch they were estimated at 5-6.10^10^ TU/ml. The behavioural sequences for the two batches are presented in Figure S8A.

For experiments with pups, P0 *Rnd2*^*flox/flox*^ mice were bilaterally injected with 1µl of GFP or GFP/Cre retroviruses on each side. The titers of GFP and GFP/Cre retroviruses were 1.10^9^ TU/ml. The behavioural sequence is presented in Figure S10A. Nine GFP and 8 GFP/Cre mice were used for the analysis.

For all behavioural tests, animals were placed in the test room 30 min before the beginning of the experiment. For all procedures, experimenters were blind to the virus injected.

### Open-field test (OF)

Mice were placed in one corner of a square open-field (50×50 cm, 200 lux). Exploratory behaviour was monitored for 10 min. Time spent in the center (35×35 cm) and the distance ratio (distance travelled in the periphery divided by the total travelled distance) were automatically measured by a video-tracking system connected to a camera (Videotrack, ViewPoint). These two parameters were used for z-open field score calculation.

### Emergence test

The emergence test was done in the same open-field but with brighter light (∼300 lux). Mice were placed in a dark cylinder (10×6.5 cm, grey PVC) in one corner of the arena. The behaviour was monitored for 5 min using the same system as in the open-field test (Videotrack, ViewPoint). Latency to emerge from the cylinder and the number of re-entries in the cylinder were automatically measured and used for z-emergence score calculation.

### Elevated Plus Maze (EPM)

Behaviour in the EPM was measured using a cross maze with two open and two closed arms (37×6 cm arms, 75 lux) placed 115 cm above the ground. Mice were placed at the centre of the cross, facing one of the open arms and the behaviour was monitored for 5 min. Time spent in open arms as well as the ratio between entries in open arms divided by entries in closed and open arms were measured using the video-tracking system (Videotrack, ViewPoint) and used for z-EPM score calculation.

### Light/Dark

The apparatus used for this test was composed of a strongly illuminated chamber (36×36 cm, ∼350 lux) and a dark chamber (36×23 cm, ∼10 lux) separated by a wall with a door (10×10 cm) allowing animals to travel freely between the two compartments. Mice were placed in the lit chamber and left exploring the apparatus for 5 min. Latency to leave the lit chamber and time spent in it were manually measured and used for z-light dark score calculation.

### Sucrose preference

In the sucrose preference test, mice were exposed in their home cage to two drinking bottles during 72 h. During the first 24 h, a habituation was performed with the two bottles filled with water. For the test phase, one of the bottles was filled with 30 mL of 4% sucrose water while the second bottle was filled with water. Bottles were placed side by side for 24 h, then for the last 24 h positions were reversed. Sucrose preference ratio was calculated as follows and used for z-sucrose preference score calculation:

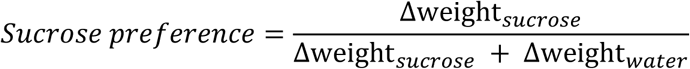

### Forced swim test (FST)

The FST device consists of a glass cylinder 16 cm in diameter and 25 cm high, filled with water to a height of 15 cm and placed in a strongly illuminated room (∼350 lux). Water was maintained at ∼26°C. Mice were placed in the apparatus and behaviour was monitored for 6 min. Latency to immobility and total immobility time in the last 4 min of the test were measured and used for z-FST score calculation.

### Morris Water Maze (MWM)

The apparatus was a white circular swimming pool (150 cm in diameter and 60 cm deep) located in a room with various distal visual cues (80 lux). The pool was filled with water maintained at 19°C and made opaque by the addition of a non-toxic white cosmetic adjuvant. The escape platform (14 cm diameter) was hidden underwater so that its top surface was 0.5 cm below the surface of the water. In this task, mice were required to locate the hidden platform using distal cues and mice behaviour was monitored using a camera and a video-tracking system (Videotrack, ViewPoint). For the entire training period, the platform stayed in the same position and mice were tested with variable random start positions (Fig. S8B). Mice were trained in 3 trials a day, each trial being separated by a 5 min interval. A trial was terminated when the animal climbed onto the platform. Mice that failed to find the platform within the 60 sec cut-off time were placed onto the platform by the experimenter and had to stay there for 15 sec before being placed back in their home cage. The releasing point (starting point) differed for each trial and different sequences of releasing points were used day to day. Three days after the last training trial, the hidden platform was removed and the memory for the platform location was assessed during a probe test (Fig. S8B). During this test, mice were allowed to freely swim in the water maze for 60 sec and performances were assessed by time spent in the target quadrant where the platform was located.

### Contextual fear conditioning

Mice were exposed to contextual fear conditioning in two distinct conditioning contexts that shared features in order to test their ability to discriminate these contexts. Conditioning was done in a transparent plexiglass cage (26×25×17 cm) allowing access to visual cues of the environment with a floor composed of 42 stainless steel rods, separated by 3 mm, which were wired to a shock generator and scrambler (Imetronic). In context A, the cage was illuminated (90 lux), the testing room was strongly illuminated (250 lux) and visual cues were placed on the walls of the room. The cage was cleaned between each mouse with a 70% ethanol solution and the experimenter wore latex gloves. For context B, the testing room was lit with a low light (50 lux) and the cage was not illuminated. Dark panels were placed proximal to the cage, on which visual cues different from those used in context A were placed. Furthermore, a plastic boat containing used litter was added under the steel bars of the cage and lemon scent was added in the cage. Between each mouse, the cage was cleaned with 30% acetic acid solution and the experimenter wore nitrile gloves.

For the conditioning phase, mice were individually transported from the resting room to the experimental room and placed in the context A conditioning chamber. After 180, 240 and 300 sec, they received a single footshock (0,7 mA, 1 sec) and remained in the chamber for 1 min (Fig. S8E) after the last shock before being transported back to their housing room. On the following day mice were tested for context discrimination. In this goal, mice were again individually transported to the experimental room and placed in the context A conditioning chamber for 5 min without delivery of a footshock. Three hours later, mice were then exposed to the context B conditioning chamber for 5 min, again without any footshock (Fig. S8E). During the conditioning phase and the testing phase, mice behaviour was monitored with a video-tracking system (Freezing, Imétronic). Freezing behaviour, defined as a behavioural immobility except for breathing movements, was measured. Discrimination of contexts was evaluated comparing the immobility time in each context, a discrimination ratio was calculated as follows:

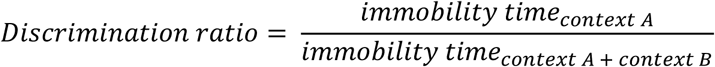

### Criteria of exclusion

Most tests used in this study rely on exploration and locomotor activity, therefore total distance travelled in these tests was used as an index of locomotor activity. Although no animal was excluded on the basis of the total travelled distance, several other criteria were determined to exclude mice that did not follow the rules of the designed tests. For example mice that did not swim and instantly started floating during the MWM and the FST would be excluded for the final analysis of performances, as well as mice that did not leave the cylinder in the emergence test. In the EPM, mice that stayed 240 sec in the centre or less than 1 sec in open or closed arms were excluded from analysis, as well as mice that explored less than 2 different arms. For behavioural experiments with adults, no animal was removed from the analysis after euthanasia because all of them showed a significant number of GFP+ cells on both side. For behavioural experiments with pups, only animals with GFP+ cells on both sides were kept for the analysis.

### Z-score calculation

Z-scores are mean-normalization of the results and allow for comparison of related data across this study. They indicate how many standard deviations (σ) an observation (X) is above or below the mean of a control group (µ) and were calculated as previously described (Guilloux et al., 2011):

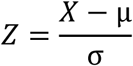

Z-score values were calculated for test parameters measuring emotionality. The directionality of scores was adjusted so that increased score values reflect increased dimensionality (anxiety or depression-related behaviour). For instance, decreased time spent in open arms in the EPM or decreased time spent in the centre of the open-field test, compared to control, were converted into positive standard deviation changes compared to group means indicating increased anxiety-related behaviour. On the other hand, increased immobility time in the FST for example, is directly related to increased depression-like behaviour and was not converted.

As an example, z-score in the open-field (Z_OF_) was calculated for each animal, using normalization of “time in the center” (TC) and “distance in periphery/total distance ratio” (DR) values.

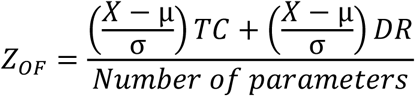

Similarly, z-scores were calculated for the emergence test (Z_emergence_), the EPM (Z_EPM_) and the light/dark test (Z_L/D_). z values obtained for each test were then averaged to obtain a single z-anxiety score:

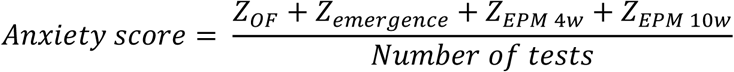

The same calculations were done with the z-scores of the sucrose preference test (Z_SP_) and the FST (Z_FST_), and an individual “depression score” was calculated as for the z-anxiety score.

### Statistical analysis

Statistical analyses were performed using the GraphPad Prism software. No statistical methods were used to predetermine sample sizes, but our sample sizes are similar to those generally employed in the field. Results are presented as mean ± s.e.m. (standard error of the mean). The statistical test used for each experiment and sample size (n) are indicated in the corresponding figure legend or in the Methods. Statistical values are mentioned in the text.

## ACKNOWLEDGMENTS

This work was supported by INSERM (to D.N.A. and E.P.), ANR (ANR-19-CE16-0014-01 to E.P.), University of Bordeaux, Marie Curie Actions (Intra European Fellowship to D.N.A. for E.P.), the Wellcome Trust (project grant 086947/Z/08/Z to F.G.). T.K. was supported by a MESR (Ministère de l’Enseignement Supérieur et de la Recherche) fellowship and by a LabEX BRAIN (Bordeaux Region Aquitaine Initiative for Neuroscience) PhD extension grant, A.G. was supported by a FRM (Fondation pour la Recherche Medicale) grant (ING20121226343 to D.N.A.) and LabEX BRAIN grant (to D.N.A.). Work in F.G.’s lab is supported by the Francis Crick Institute, which receives its funding from Cancer Research UK (FC0010089), the UK Medical Research Council (FC0010089) and the Wellcome Trust (FC0010089). We are grateful to Sarah Hart-Jonson and the Biological Services at the National Institute for Medical Research (NIMR), UK for generating the *Rnd2*^*flox/flox*^ mutant mouse line. We thank Dr Vidya Ramesh and Dr Roberta Azzarelli for assistance in breeding and genotyping this mouse line in NIMR. We thank Dr Fred Gage for providing CAG–GFP, CAG-GFP/Cre, CAG-RFP and CAG-GFP-T2A-CreER^T2^ retroviral vectors, Dr Benedikt Berninger for CAG-IRES-DsRed and CAG-Neurog2-IRES-DsRed retroviral vectors and Dr Dieter Chichung Lie for providing the 293GPG cell line. We gratefully acknowledge Cedric Dupuy and Fiona Corailler for animal care in the Neurocentre Magendie. We thank Dr Francis Chaouloff and Dr Giovanni Marsicano for lending the light/dark apparatus and the electrophysiological setup respectively. This work benefited from the support of the Microdissection Laser Capture facility funded by INSERM, LabEX BRAIN ANR-10-LABX-43 and FRM DGE20061007758, the Transcriptomic facility funded by INSERM and LabEX BRAIN ANR-10-LABX-43, the Biochemistry and Biophysics Facility of the Bordeaux Neurocampus funded by the LabEX BRAIN ANR-10-LABX-43 and the Animal Housing and Genotyping facilities funded by INSERM and LabEX BRAIN ANR-10-LABX-43. Confocal microscopy was done in the Bordeaux Imaging Center (BIC), a service unit of the CNRS-INSERM and Bordeaux University, member of the national infrastructure France BioImaging supported by the French National Research Agency (ANR-10-INBS-04).

## AUTHOR CONTRIBUTIONS

E.P., T.K. and D.N.A. conceived the experiments, analyzed the data and wrote the manuscript. T.K., together with E.P., carried out most of the experiments. F.F. prepared DNA and produced retroviruses together with A.G. M.M. and H.D. performed microdissection analyses. G.T. performed the electrophysiological experiments. T.L.-L. and H.D. performed PCR analyses. M.K. and D.N.A. designed behavioural experiments. M.B. performed tests in the MWM. J.I.H. designed the *Rnd2* conditional mutant allele together with F.G. who then generated the *Rnd2* conditional mutant mouse. D.G. supervised the breeding and genotyping of *Rnd2*^*flox/flox*^ mice.

## DECLARATION OF INTERESTS

The authors declare no conflict of interest.

## Notes

### Competing Interest Statement

The authors have declared no competing interest.

