## Supplementary material for "The atypical Rho GTPase Rnd2 is critical for dentate granule neuron development and anxiety-like behavior during adult but not neonatal neurogenesis"

### SUPPLEMENTARY FIGURE LEGENDS

**Supplementary Figure 1: *Rnd1* and *Rnd3* expressions in the mouse DG.**

(**A**) Sequential images of the hippocampus showing the region (SGZ and GCL) microdissected by laser capture from cresyl violet-stained sections. (**B**) Analysis by real-time PCR of *Rnd1* and *Rnd3* mRNA expressions in microdissected DG (SGZ and GCL) at different ages. Data are presented as fold change compared with the expression level at P7 ± s.e.m (n = 4 mice per time point). P, postnatal day; W, weeks. (**C**) Distribution of *Rnd1* and *Rnd3* transcripts in the adult hippocampus (12-week old mouse). **(D**) Analysis by real-time PCR of *Rnd1* and *Rnd3* mRNA expressions along the septo-temporal axis in the adult (12-week old) microdissected DG. Data are presented as fold change compared with the expression level at the anteroposterior coordinate -1.34 from the Bregma ± s.e.m (n = 5 mice per time point).

Scale bar represents 300 µm (A, C).

**Supplementary Figure 2: *Rnd2* deletion in adult-born DGNs using *Rnd2*$^{flox/flox}$ mice and Cre expressing retrovirus.**

(**A**) *Rnd2* conditional mutant allele. This allele contains one loxP site between the exon 1 and 2 and a second loxP site after the last exon. A neomycin (Neo) selection gene flanked by flippase recognition target (FRT) sites was inserted in 3' of *Rnd2*. Following transmission of the mutation to the germline, the Neo gene was excised, giving rise to the *Rnd2*$^{flox}$ allele. Upon Cre recombination, exons 2 to 5 of *Rnd2* are deleted. (**B**) Analysis 7 days post-retroviral injection of *Rnd1*, *Rnd2* and *Rnd3* transcripts by quantitative RT-PCR in micro-dissected GFP+ cells. Graphs show expression levels normalized to housekeeping genes and relative to *Rnd* expression in control condition (GFP). (n=9 for GFP condition, n=8 for GFP/Cre condition; 500-800 cells per animal). Mean ± s.e.m., Unpaired two-tailed Student's t-test; ***p < 0.001.

**Supplementary Figure 3: *Rnd2* deletion does not impact the proliferation and neuronal differentiation of adult-born DGNs.**

(**A**) Immunostainings for GFP and Ki67 in the DG 3 days after GFP (control) or GFP/Cre (*Rnd2* deletion) retrovirus injection. TOTO labels nuclei (**B**) Quantification of the percentage of transduced cells that are Ki67+ at 3 and 7 days post-injetion (dpi). Mean ± s.e.m (n = 4-6 mice). (**C**) Immunostainings for GFP and doublecortin (DCX) in the DG 14 days after virus

injection. (**B**) Quantification of the percentage of transduced cells that are DCX+ at 7, 14 and 21 dpi. Mean ± s.e.m (n = 4-7 mice).
Scale bars represent 20 μm (A, C).

**Supplementary Figure 4: The exacerbated cell death is specific to *Rnd2* suppression.**
(**A**) The retroviral co-injection strategy was used in C57Bl6/J mice. The graph shows the survival rate of only RFP+ cells and total GFP+ cells. Mean ± s.e.m.; paired two-tailed Student's t-test; (n = 8 mice for each time point). (**B**) Experimental design for survival rescue experiments. Double stained (DS) cells were quantified 3 and 21 days after injection of a mixture of retroviruses into the DG of adult *Rnd2$^{flox/flox}$* mice. (**C**) Survival rate of DS cells at 21 days post-injection (dpi). The number of DS cells at 21 dpi was normalized to the number obtained for the corresponding viral mixture at 3 dpi. Mean ± s.e.m.; one way ANOVA followed by Dunnett's multiple comparison test; *p < 0.05 compared to Rnd2 rescue (n = 8-10 mice).

**Supplementary Figure 5: *Rnd2* deletion does not have a major impact on the dendritic spines of adult-born DGNs.**
(**A**) Representative images of control and *Rnd2*-deleted dendritic segments at 21 and 28 dpi. (**B**, **C**, **D**) Quantification of the total spine density (**B**) and the density of the different spine types at 21 dpi (**C**) and 28 dpi (**D**). Mean ± s.e.m.; paired two-tailed Student's t-test; *p < 0.05 (n = 6 at 21 dpi and n = 4 mice at 28 dpi, a minimum of 3 cells were analyzed per animal).
Scale bar represents 5 μm (A).

**Supplementary Figure 6: *Rnd2* deletion at 28 dpi induces cell death but does affect cell positioning and morphology.**
(**A**) A retrovirus expressing GFP together with a conditionally active form of Cre recombinase, which is activated upon tamoxifen, was injected into the DG of adult *Rnd2$^{flox/flox}$* mice. Four weeks after virus injection, tamoxifen (150 mg/kg, daily for 5 days), or oil in control group, was injected and animals were sacrificed 21 days after the last injection of tamoxifen. (**B**) The relative number of GFP+ cells, the relative position in the GCL and the morphology of transduced cells were quantified 21 days after the last injection of tamoxifen. Mean ± s.e.m., Unpaired two-tailed Student's t-test; *p<0.05 (n=5-8 animals per group).

**Supplementary Figure 7: *Rnd2* deletion does not impact the membrane properties of adult-born DGNs.**

(**A**) Images of adult newborn neurons transduced with GFP or GFP/Cre retrovirus at 28 dpi in the DG of *Rnd2*$^{flox/flox}$ mice. Biocytin identifies neurons in which whole-cell patch-clamp recording was performed. (**B**, **C**, **D**) Quantification of the resting membrane potential (A), membrane capacitance (B) and membrane resistance (C). Mean ± s.e.m (n = 30 neurons from 4 mice in GFP group and n = 17 neurons from 8 mice in GFP/Cre group).

Scale bar represents 20 μm (A).

**Supplementary Figure 8: *Rnd2* suppression in adult-born DGNs, using a retroviral approach, does not perturb hippocampal-dependent memory.**

(**A**) Experimental design and diagram of the behavioral task sequences. (**B**) Experimental design used in the Morris water maze. NW, quadrant north west; NE, quadrant north east; SW, quadrant south west; SE, quadrant south east. (**C**) Latency to reach the hidden platform using variable start positions, 5 weeks after GFP or GFP/Cre retrovirus injection. (**D**) Time spent in the different quadrants during the probe test. (**E**) Experimental design used for contextual fear conditioning and context discrimination. (**F, G**) Contextual fear conditioning assessed by the percentage of freezing when mice were re-exposed (**F**) 24 hours or (**G**) 5 weeks later to the conditioning context (context A) and to a similar context (context B). The percentage of freezing for the baseline was measured during the first three minutes before the first shock. Mean ± s.e.m., two way ANOVA; $^{**}p<0.01$ $^{***}p<0.001$.

**Supplementary Figure 9: *Rnd2* suppression in adult-born DGNs increases anxiety-like behavior but does not affect depressive-like behavior.**

(**A**) Total travelled distance in the arena and (**B**) time spent in the center of the open field, 4 weeks after GFP or GFP/Cre retrovirus injection. (**C**) Total travelled distance, (**D**) latency to emerge from the cylinder and (**E**) number of re-entries in the cylinder in the emergence test, 4 weeks after virus infusion. (**F**) Sucrose preference (% of total intake) and (**G**) total intake 9 weeks after GFP or GFP/Cre virus injection into *Rnd2*$^{flox/flox}$ mice. (**H**) Latency to first immobility and (**I**) total immobility time in the forced-swim test, 11 weeks after retrovirus injection. (**J**) Final z-depression score after averaging z-score values of individual test. (**K-N**) Quantification of the number of transduced (GFP+) cells at the end of the behavioral sequences. The number of GFP+ cells along the septo-temporal axis (K, M) is shown as well as the total number per mouse (L, N) for the first (K, L) and second (M, N) batch. All data are presented as the mean ± s.e.m., Unpaired two-tailed Student's t-test; $^{**}p<0.01$, $^{***}p<0.001$. Batch 1 (GFP n=13; GFP/Cre n=14), Batch 2 (GFP n=11; GFP/Cre n=9).

**Supplementary Figure 10: *Rnd2* suppression in neonatally-born DGNs does not affect anxiety-like behavior in adult mice.**
(**A**) Experimental design and diagram of the behavioral task sequence. The behavioral tasks to measure anxiety-like behavior were performed 16 weeks after GFP or GFP/Cre retrovirus injection into the DG of P0 $Rnd2^{flox/flox}$ pups. (**B**) Total travelled distance in the arena and (**C**) time spent in the center of the open field. (**D**) Total travelled distance, (**E**) latency to emerge from the cylinder and (**F**) number of re-entries in the cylinder in the emergence test. (**G-H**) Quantification of the number of GFP+ cells at the end of the behavioral sequences. The number of GFP+ cells along the septo-temporal axis (H) is shown as well as the total number per mouse (G). Mean ± s.e.m., Unpaired two-tailed Student's t-test, GFP n=9 and GFP/Cre n=8 mice.

FIGURE S1

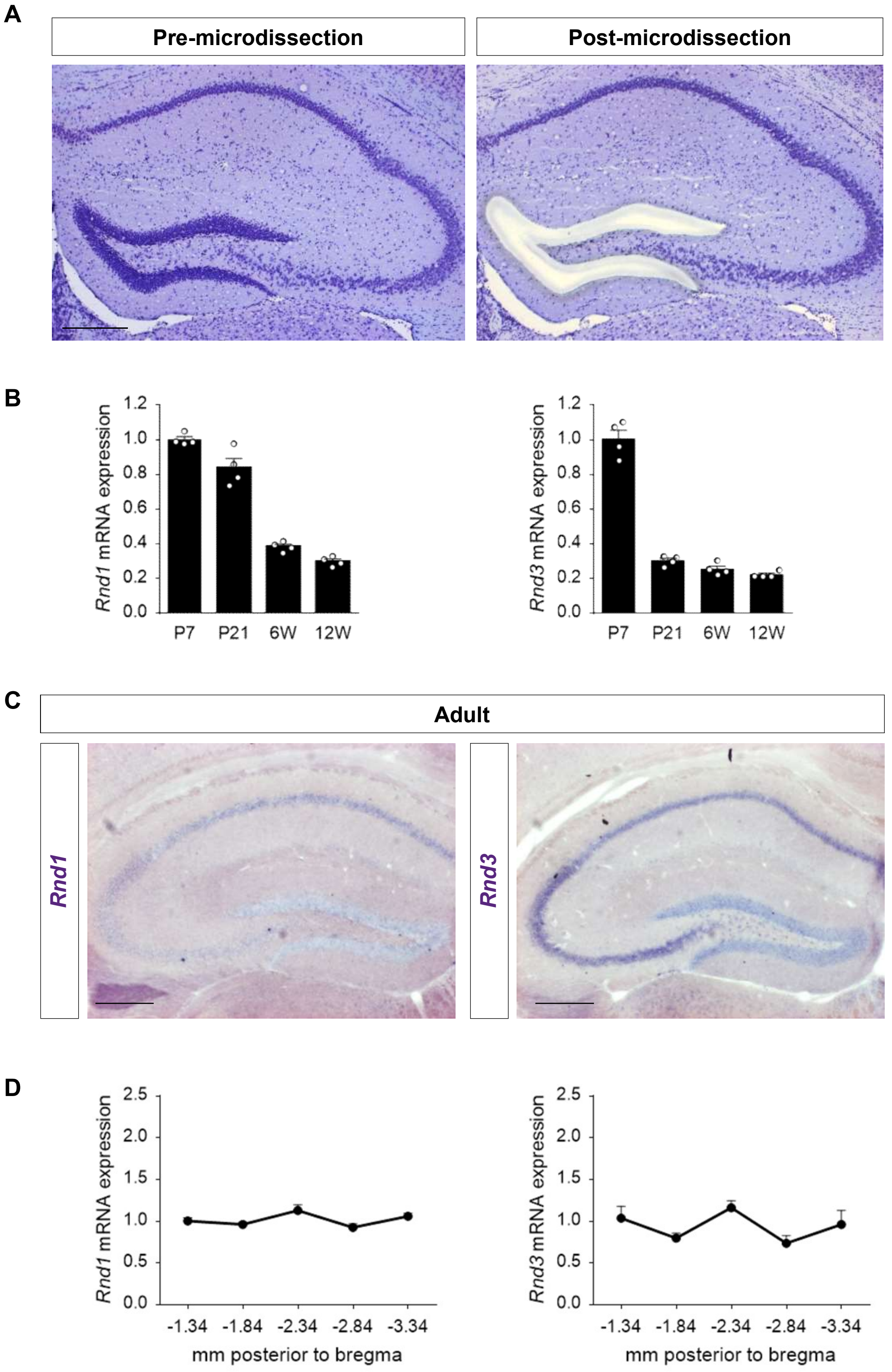
A
Pre-microdissection
Post-microdissection
B
Rnd1 mRNA expression
1.2
1.0
0.8
0.6
0.4
0.2
0.0
P7
P21
6W
12W
Rnd3 mRNA expression
C
Adult
Rnd1
Rnd3
D
2.5
2.0
1.5
1.0
0.5
0.0
-1.34
-1.84
-2.34
-2.84
-3.34
mm posterior to bregma

**FIGURE S2**

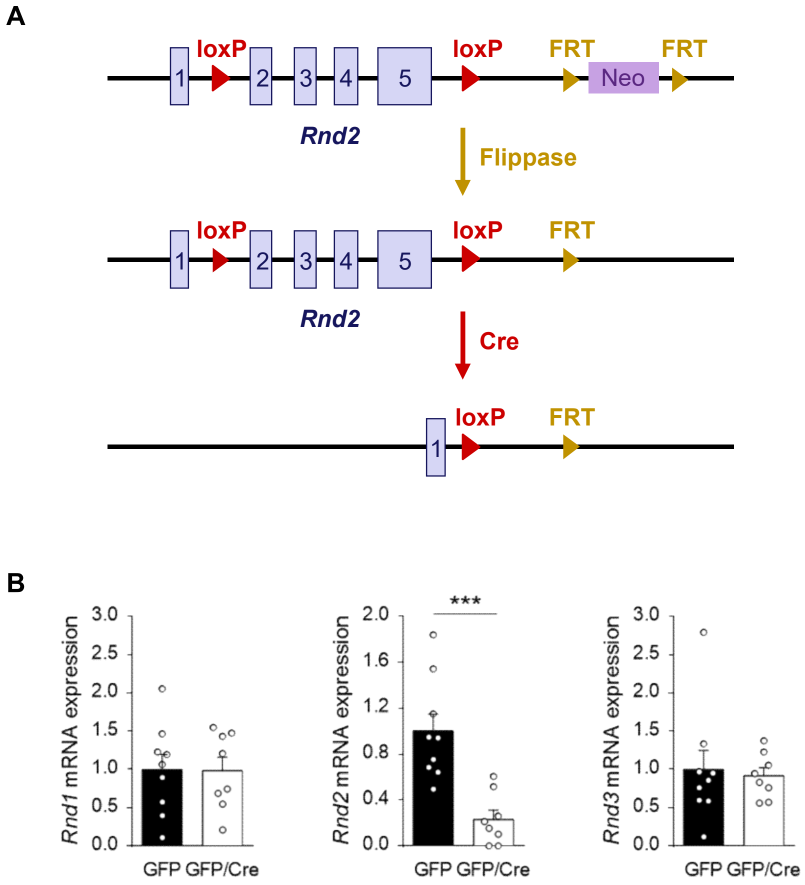
A
1
loxP
2
3
4
5
loxP
FRT
Neo
FRT
Rnd2
Flippase
1
loxP
2
3
4
5
loxP
FRT
Rnd2
Cre
1
loxP
FRT
B
Rnd1 mRNA expression
3.0
2.5
2.0
1.5
1.0
0.5
0.0
GFP GFP/Cre
***
Rnd2 mRNA expression
2.0
1.6
1.2
0.8
0.4
0.0
GFP GFP/Cre
Rnd3 mRNA expression
3.0
2.5
2.0
1.5
1.0
0.5
0.0
GFP GFP/Cre

FIGURE S3

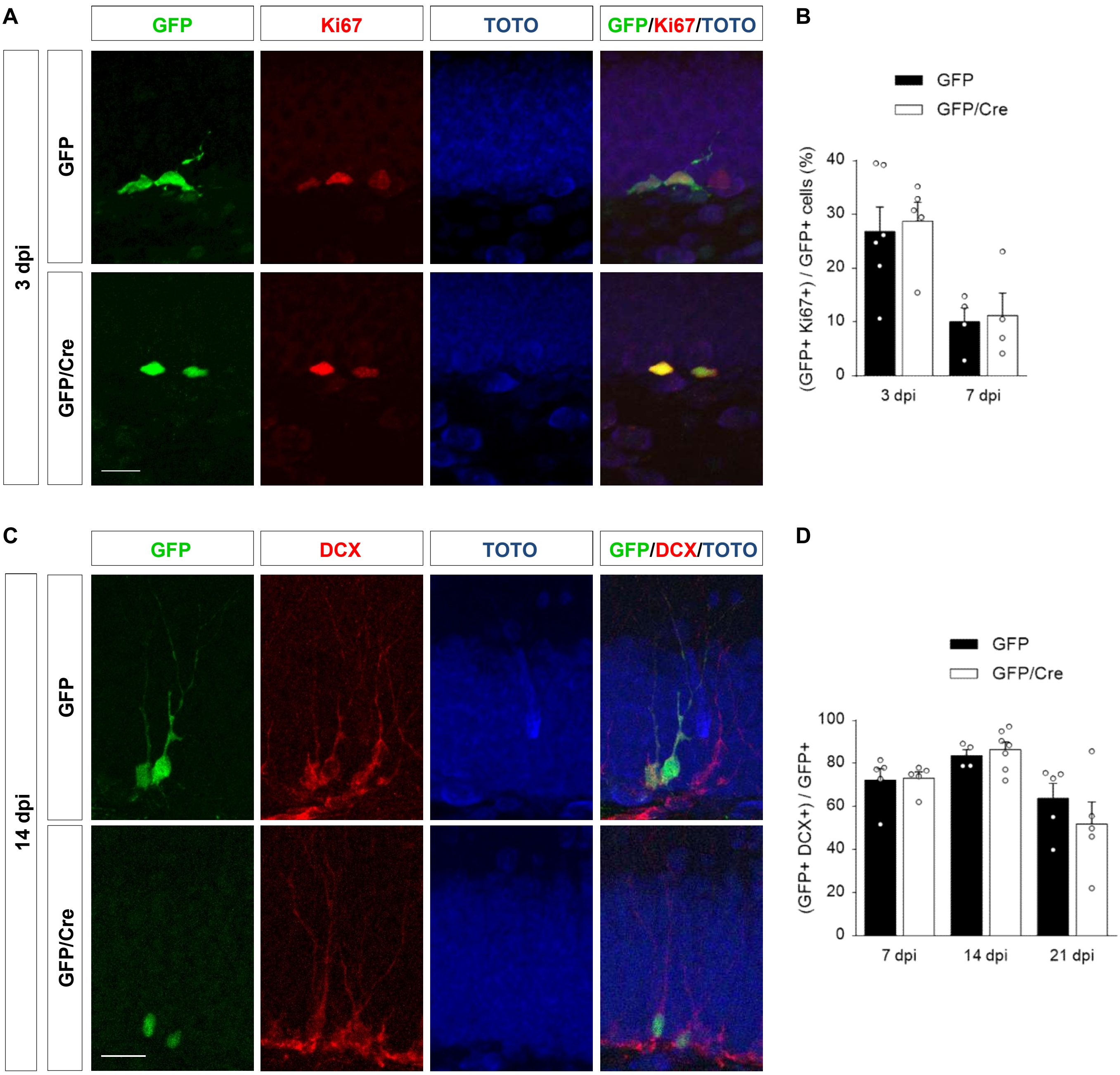

A
GFP
Ki67
TOTO
GFP/Ki67/TOTO
3 dpi
GFP
GFP/Cre
B
GFP
GFP/Cre
(GFP+ Ki67+) / GFP+ cells (%)
40
30
20
10
0
3 dpi
7 dpi
C
GFP
DCX
TOTO
GFP/DCX/TOTO
14 dpi
GFP
GFP/Cre
D
GFP
GFP/Cre
(GFP+ DCX+) / GFP+
100
80
60
40
20
0
7 dpi
14 dpi
21 dpi

**FIGURE S4**

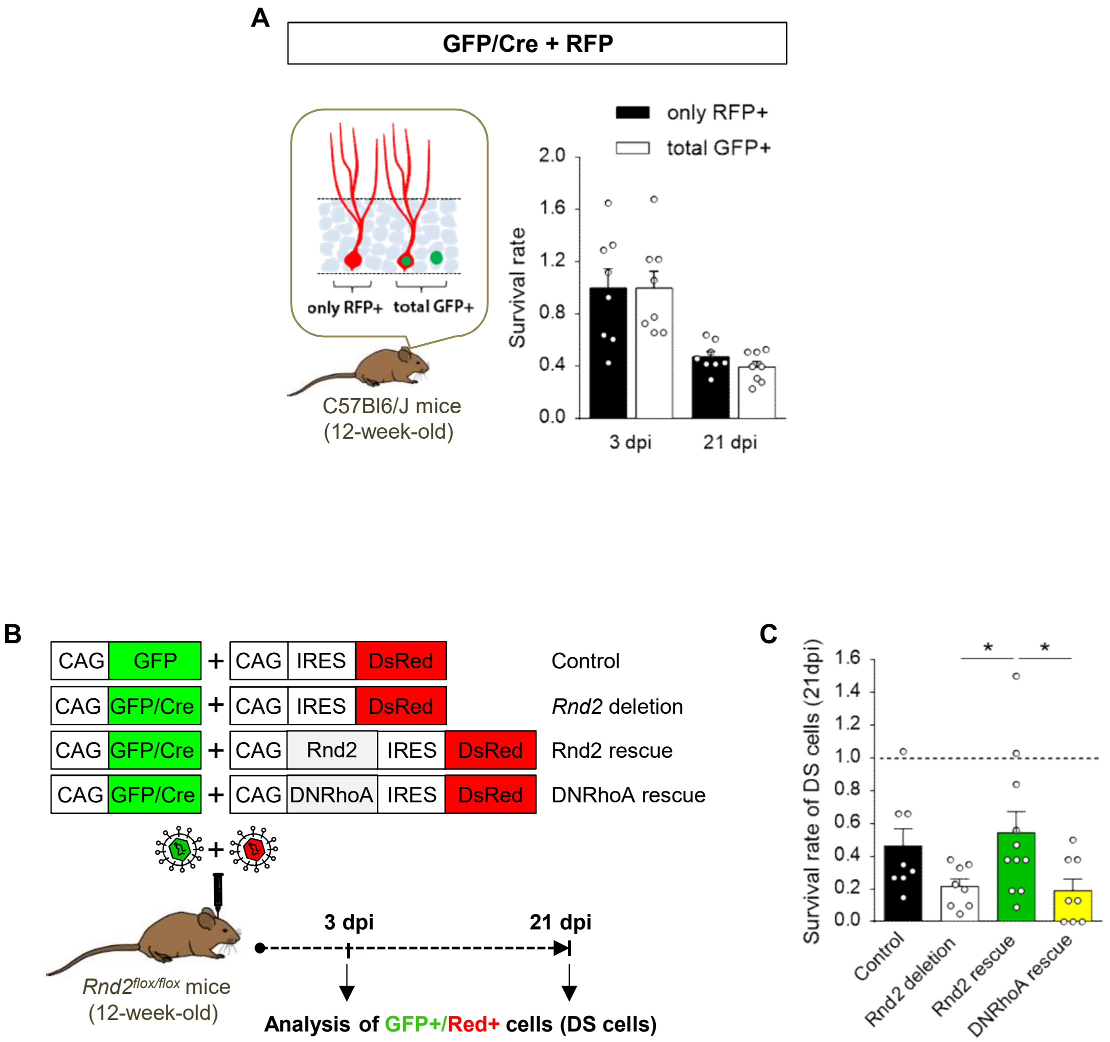

A
GFP/Cre + RFP
only RFP+
total GFP+
C57Bl6/J mice
(12-week-old)
only RFP+
total GFP+
Survival rate
2.0
1.6
1.2
0.8
0.4
0.0
3 dpi
21 dpi
B
CAG GFP + CAG IRES DsRed
Control
CAG GFP/Cre + CAG IRES DsRed
Rnd2 deletion
CAG GFP/Cre + CAG Rnd2 IRES DsRed
Rnd2 rescue
CAG GFP/Cre + CAG DNRhoA IRES DsRed
DNRhoA rescue
Rnd2flox/flox mice
(12-week-old)
3 dpi
21 dpi
Analysis of GFP+/Red+ cells (DS cells)
C
Survival rate of DS cells (21dpi)
1.6
1.4
1.2
1.0
0.8
0.6
0.4
0.2
0.0
*
*
Control
Rnd2 deletion
Rnd2 rescue
DNRhoA rescue

FIGURE S5

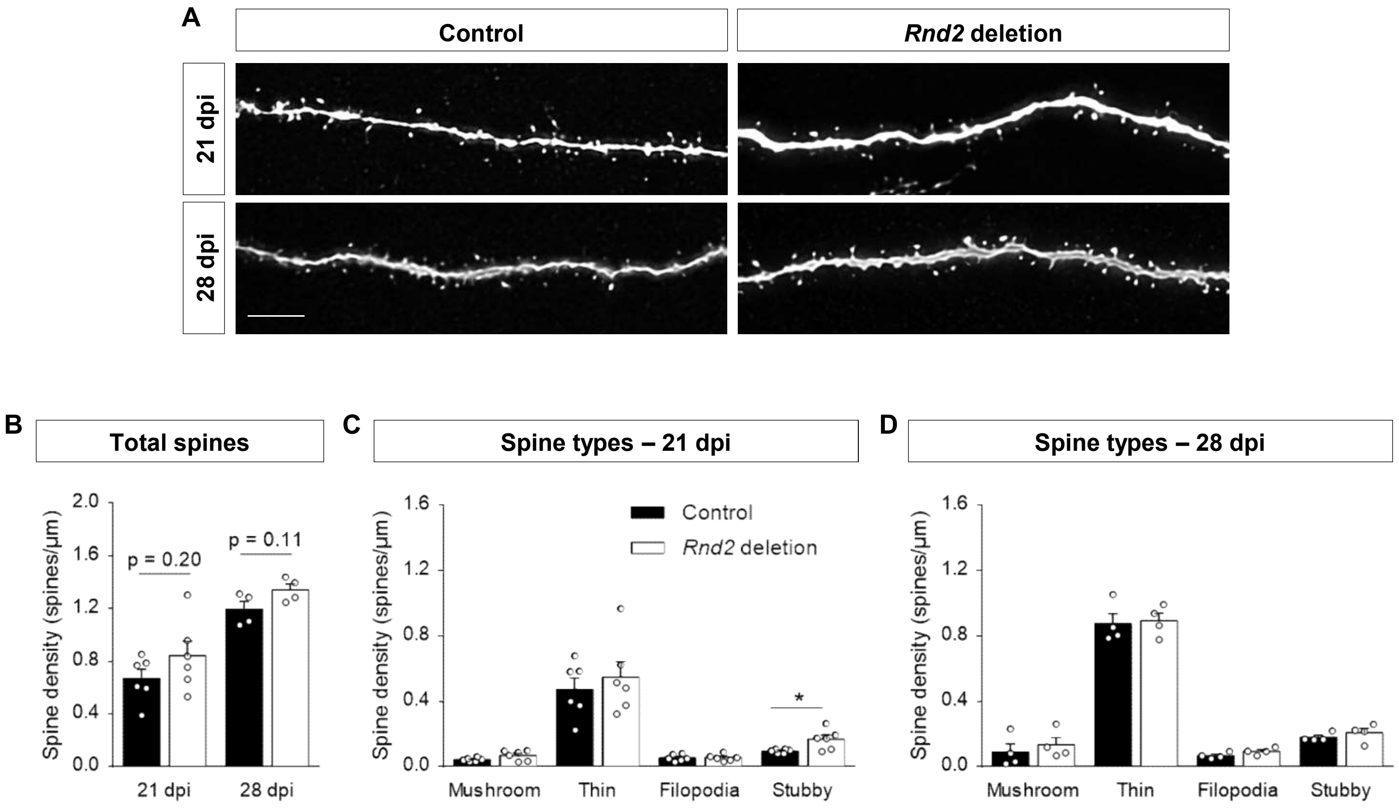
A
Control
Rnd2 deletion
21 dpi
28 dpi
B
Total spines
Spine density (spines/µm)
p = 0.20
p = 0.11
21 dpi
28 dpi
C
Spine types – 21 dpi
Control
Rnd2 deletion
*
Mushroom
Thin
Filopodia
Stubby
D
Spine types – 28 dpi
Mushroom
Thin
Filopodia
Stubby

**FIGURE S6**

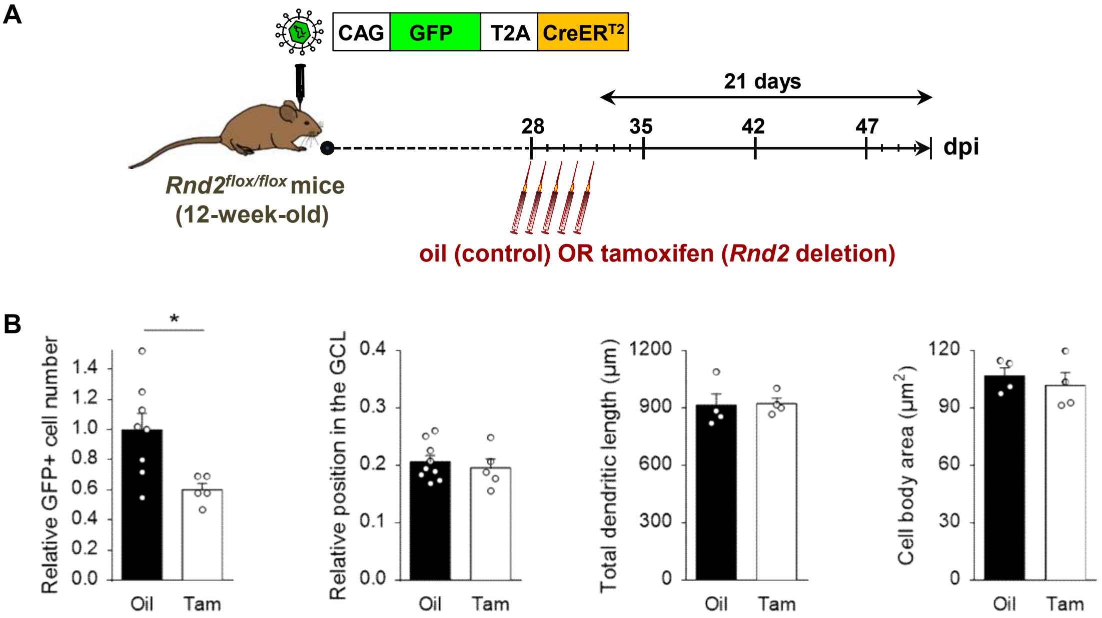
A
CAG
GFP
T2A
CreER$^{T2}$
21 days
28
35
42
47
dpi
*Rnd2$^{flox/flox}$* mice
(12-week-old)
oil (control) OR tamoxifen (*Rnd2* deletion)
B
Relative GFP+ cell number
*
Relative position in the GCL
Total dendritic length (µm)
Cell body area (µm$^2$)
Oil
Tam

**FIGURE S7**

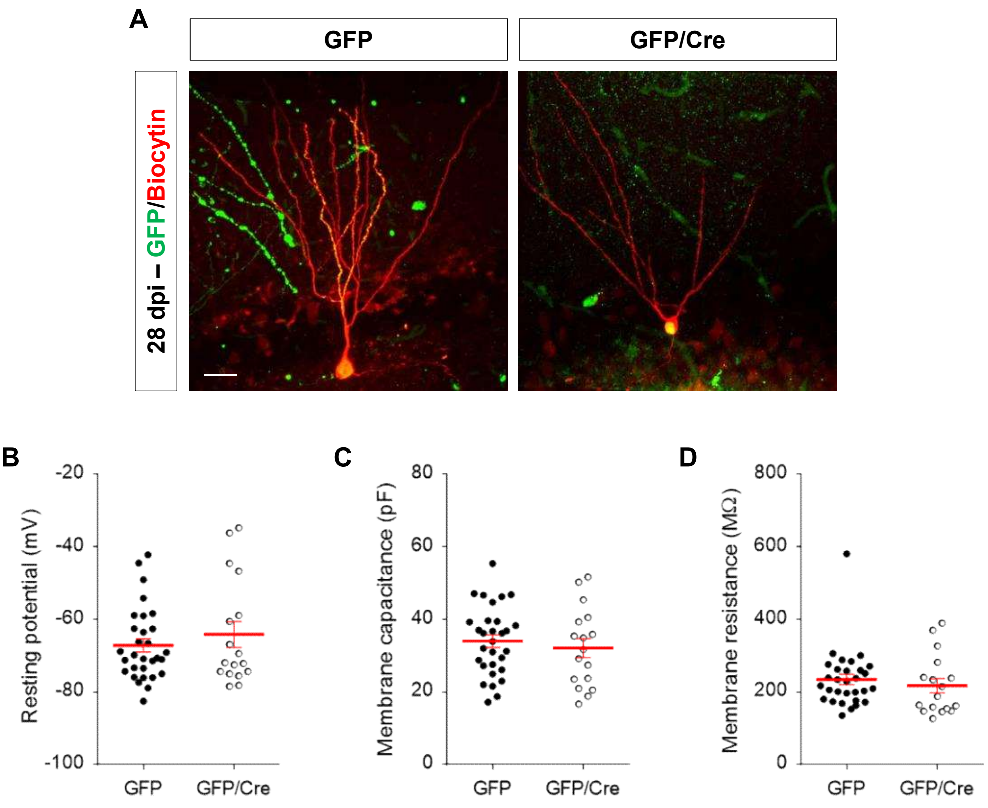
A
GFP
GFP/Cre
28 dpi – GFP/Biocytin
B
Resting potential (mV)
-20
-40
-60
-80
-100
GFP
GFP/Cre
C
Membrane capacitance (pF)
80
60
40
20
0
GFP
GFP/Cre
D
Membrane resistance (MΩ)
800
600
400
200
0
GFP
GFP/Cre

FIGURE S8

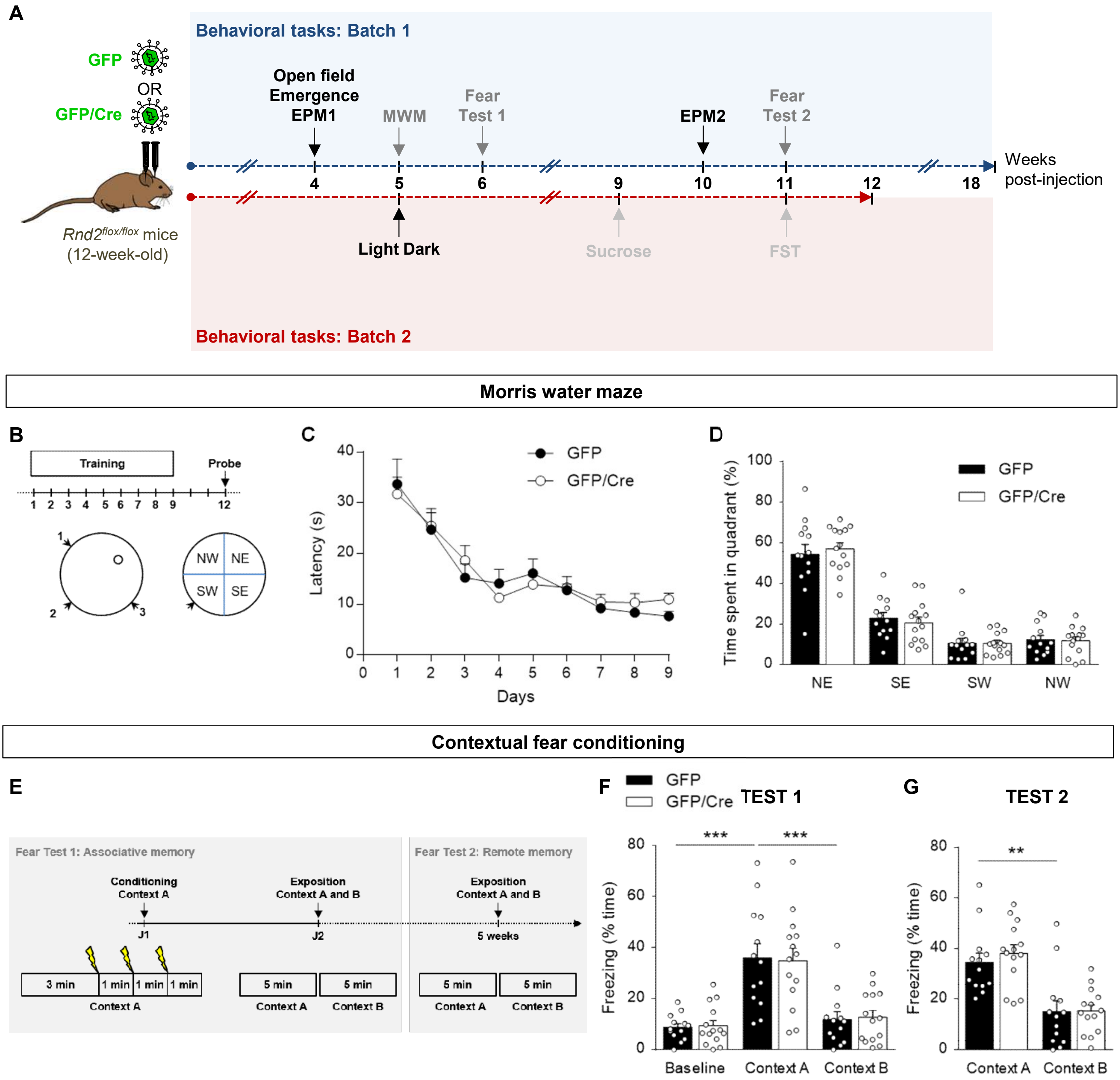
A
Behavioral tasks: Batch 1
GFP
OR
GFP/Cre
Rnd2flox/flox mice
(12-week-old)
Open field
Emergence
EPM1
MWM
Fear
Test 1
EPM2
Fear
Test 2
4
5
6
9
10
11
12
18
Weeks
post-injection
Light Dark
Sucrose
FST
Behavioral tasks: Batch 2
Morris water maze
B
Training
Probe
1 2 3 4 5 6 7 8 9 12
NW NE
SW SE
C
GFP
GFP/Cre
Latency (s)
Days
D
Time spent in quadrant (%)
NE SE SW NW
Contextual fear conditioning
E
Fear Test 1: Associative memory
Fear Test 2: Remote memory
Conditioning
Context A
Exposition
Context A and B
J1
J2
5 weeks
3 min
1 min
5 min
Context A
Context B
F
TEST 1
Freezing (% time)
Baseline
Context A
Context B
***
G
TEST 2
**

FIGURE S9

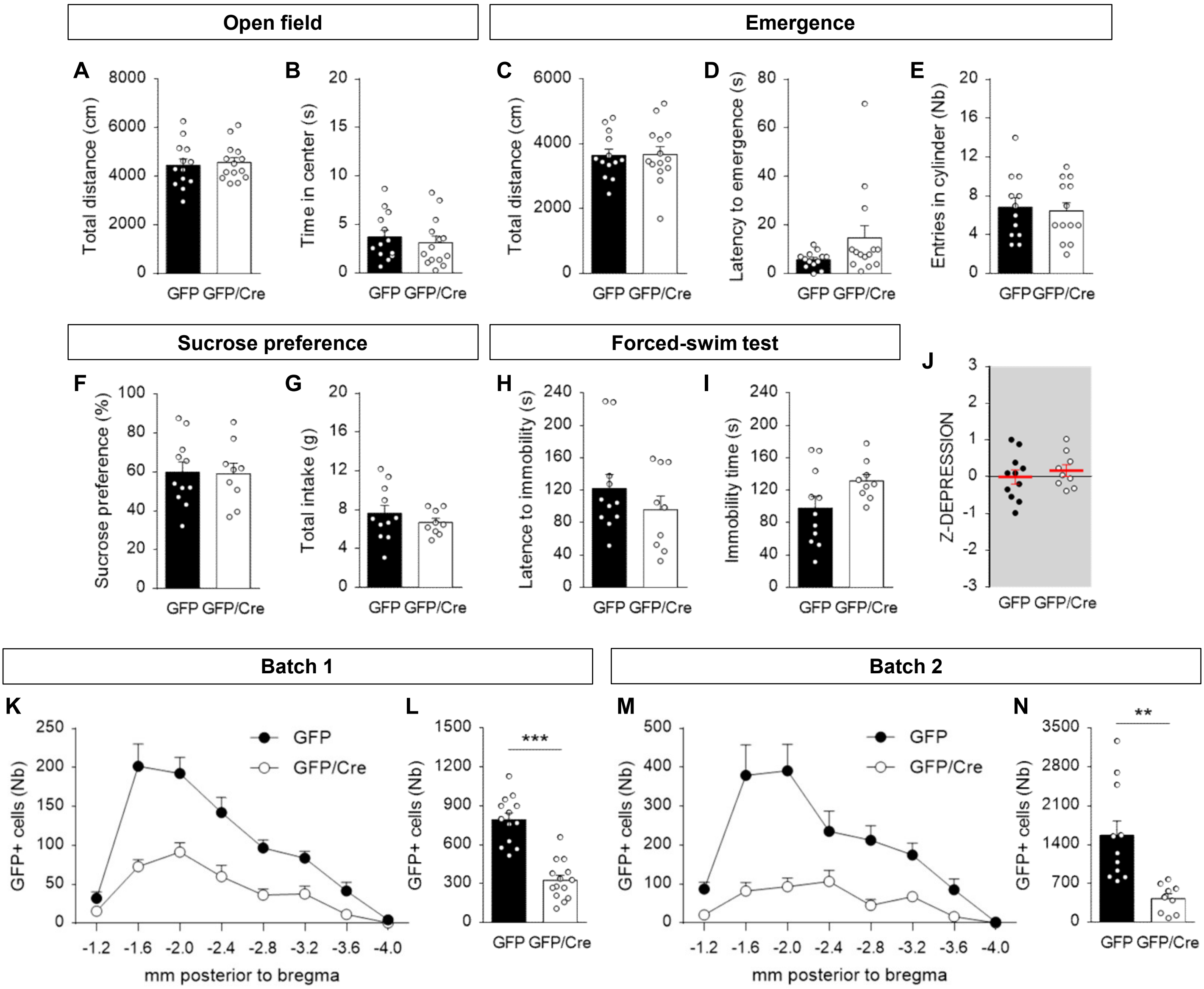

Open field
Emergence
A
Total distance (cm)
B
Time in center (s)
C
Total distance (cm)
D
Latency to emergence (s)
E
Entries in cylinder (Nb)
GFP GFP/Cre
Sucrose preference
Forced-swim test
F
Sucrose preference (%)
G
Total intake (g)
H
Latence to immobility (s)
I
Immobility time (s)
J
Z-DEPRESSION
Batch 1
Batch 2
K
GFP+ cells (Nb)
GFP
GFP/Cre
mm posterior to bregma
L
***
M
N
**

FIGURE S10

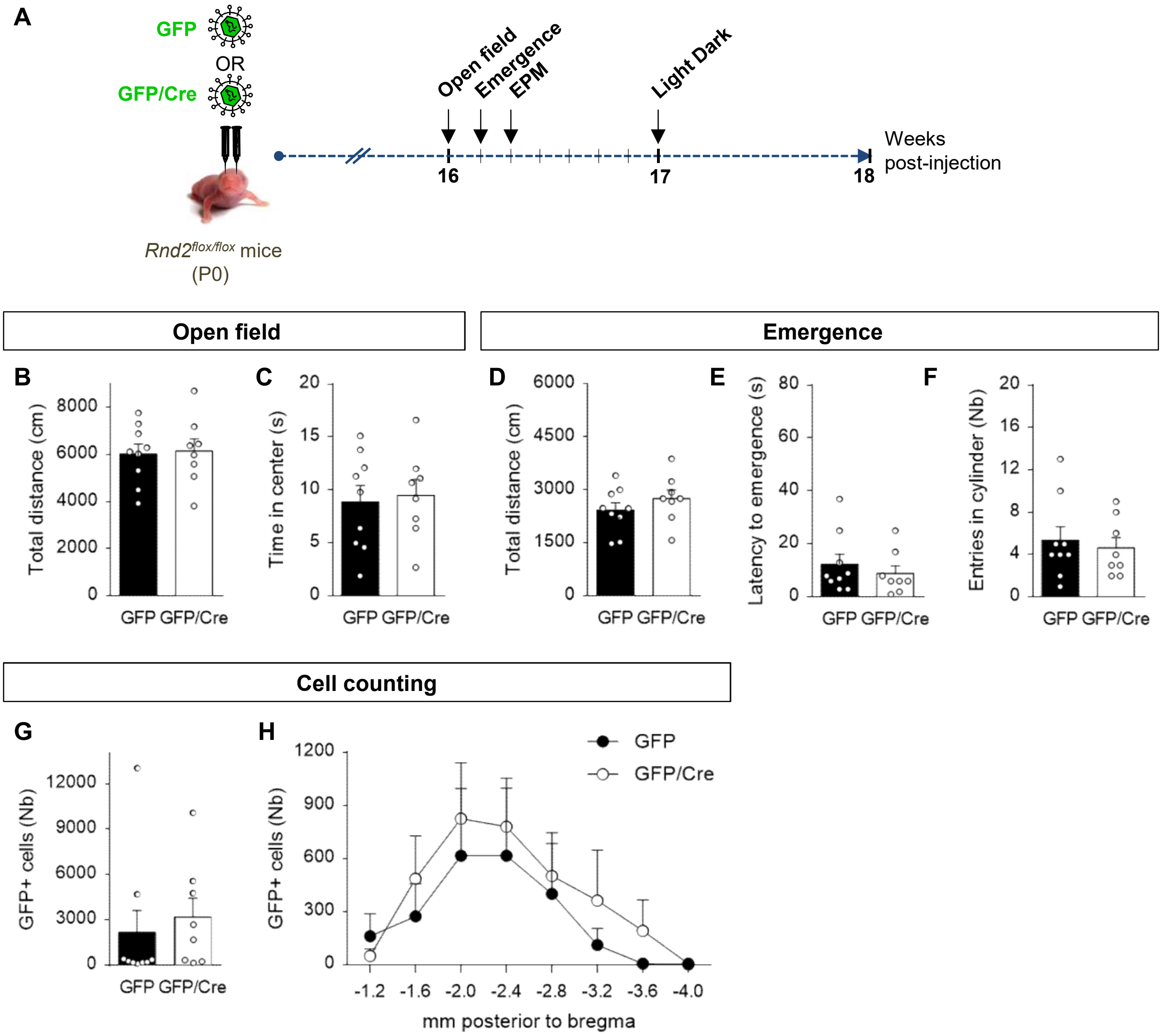

A
GFP
OR
GFP/Cre
Open field
Emergence
EPM
Light Dark
16
17
18
Weeks
post-injection
Rnd2flox/flox mice
(P0)
Open field
Emergence
B
C
D
E
F
Total distance (cm)
Time in center (s)
Total distance (cm)
Latency to emergence (s)
Entries in cylinder (Nb)
GFP GFP/Cre
Cell counting
G
H
GFP+ cells (Nb)
GFP
GFP/Cre
-1.2 -1.6 -2.0 -2.4 -2.8 -3.2 -3.6 -4.0
mm posterior to bregma
